# Whole exome sequencing and characterization of coding variation in 49,960 individuals in the UK Biobank

**DOI:** 10.1101/572347

**Authors:** Cristopher V. Van Hout, Ioanna Tachmazidou, Joshua D. Backman, Joshua X. Hoffman, Bin Ye, Ashutosh K. Pandey, Claudia Gonzaga-Jauregui, Shareef Khalid, Daren Liu, Nilanjana Banerjee, Alexander H. Li, O’Dushlaine Colm, Anthony Marcketta, Jeffrey Staples, Claudia Schurmann, Alicia Hawes, Evan Maxwell, Leland Barnard, Alexander Lopez, John Penn, Lukas Habegger, Andrew L. Blumenfeld, Ashish Yadav, Kavita Praveen, Marcus Jones, William J. Salerno, Wendy K. Chung, Ida Surakka, Cristen J. Willer, Kristian Hveem, Joseph B. Leader, David J. Carey, David H. Ledbetter, Geisinger-Regeneron DiscovEHR Collaboration, Lon Cardon, George D. Yancopoulos, Aris Economides, Giovanni Coppola, Alan R. Shuldiner, Suganthi Balasubramanian, Michael Cantor, Matthew R. Nelson, John Whittaker, Jeffrey G. Reid, Jonathan Marchini, John D. Overton, Robert A. Scott, Gonçalo Abecasis, Laura Yerges-Armstrong, Aris Baras, on behalf of the Regeneron Genetics Center

## Abstract

The UK Biobank is a prospective study of 502,543 individuals, combining extensive phenotypic and genotypic data with streamlined access for researchers around the world. Here we describe the first tranche of large-scale exome sequence data for 49,960 study participants, revealing approximately 4 million coding variants (of which ~98.4% have frequency < 1%). The data includes 231,631 predicted loss of function variants, a >10-fold increase compared to imputed sequence for the same participants. Nearly all genes (>97%) had ≥1 predicted loss of function carrier, and most genes (>69%) had ≥10 loss of function carriers. We illustrate the power of characterizing loss of function variation in this large population through association analyses across 1,741 phenotypes. In addition to replicating a range of established associations, we discover novel loss of function variants with large effects on disease traits, including *PIEZO1* on varicose veins, *COL6A1* on corneal resistance, *MEPE* on bone density, and *IQGAP2* and *GMPR* on blood cell traits. We further demonstrate the value of exome sequencing by surveying the prevalence of pathogenic variants of clinical significance in this population, finding that 2% of the population has a medically actionable variant. Additionally, we leverage the phenotypic data to characterize the relationship between rare *BRCA1* and *BRCA2* pathogenic variants and cancer risk. Exomes from the first 49,960 participants are now made accessible to the scientific community and highlight the promise offered by genomic sequencing in large-scale population-based studies.

## INTRODUCTION

The UK Biobank (UKB) is a prospective population-based study of over 500,000 individuals with extensive and readily accessible phenotypic and genetic data^1^. The release of genome-wide genotyping array data^2^ for study participants has accelerated genomic discovery through association studies, and enabled advances in population genetic analyses, the exploration of genetic overlap between traits, and Mendelian randomization studies^3,4^. While array data in combination with genotype imputation capture the spectrum of common genetic variants, rare variation that is more likely to modify protein sequences and have large phenotypic consequences is less well captured through these approaches.

Here, we extend the UKB resource with the first tranche of whole exome sequencing (WES) for 49,960 UK Biobank participants, generated by the Regeneron Genetics Center, as part of a collaboration with GlaxoSmithKline. These data are available to approved researchers through the UKB Data Showcase (see **URLs**). Exome sequencing allows direct assessment of protein-altering variants, whose functional consequences are more readily interpretable than non-coding variants, providing a clearer path towards mechanistic and therapeutic insights, as well as potential utility in therapeutic target discovery and validation^5–8^ and in precision medicine^9,10^. Here, we provide an overview of sequence variation in UKB exomes, review predicted damaging variants and their consequences in the general population and perform comprehensive loss of function (LOF) burden testing with 1,741 phenotypes, illustrating its utility in studies of common and rare phenotypes with a focus on deleterious coding variation.

## RESULTS

### Demographics and Clinical Characteristics of Sequenced Participants

A total of 50,000 participants were selected, prioritizing individuals with more complete phenotype data: those with whole body MRI imaging data from the UK Biobank Imaging Study, enhanced baseline measurements, hospital episode statistics (HES), and/or linked primary care records. Additionally, we selected one disease area for enrichment, including individuals with admission to hospital with a primary diagnosis of asthma (ICD10 codes J45 or J46). This resulted in 8,250 participants with asthma, or ~16% among sequenced participants, compared to ~13% among all 502,543 UKB participants (Table 1 and Ext. Data HESinWESvs500k_V1.xlsx). During data generation, samples from 40 participants were excluded due to failed quality control measures or participant withdrawal (Supplemental Methods), resulting in a final set of 49,960 individuals. The sequenced participants are representative of the overall 502,543 UKB participants (Table 1) for age, sex and ancestry. Due to the ascertainment strategy, sequenced participants were more likely to have HES diagnosis codes (84.2% among WES vs. 78.0% overall), were enriched for asthmatics, and many enhanced physical measures (eye measures, hearing test, electrocardiogram, Table 1). The exome sequenced participants did not differ from all participants in the median number of primary and secondary ICD10 codes. The sequenced subset includes 194 parent-offspring pairs, including 26 mother-father-child trios, 613 full-sibling pairs, 1 monozygotic twin pair and 195 second-degree genetically determined relationships^11^ based on identity-by-descent (IBD) estimates between pairs of individuals in UKB WES is included in Sup. Figure 1.

**Table 1.** Clinical characteristics in whole exome sequenced and all UK Biobank participants. Demographics and clinical characteristics of UKB 50K sequenced participants and overall 500K participants. See Supplemental Methods for definition of UKB clinical phenotype definitions. Unless otherwise noted, values are expressed as median (1^st^ and 3^rd^ quartile). ^a^The number of samples with exome sequencing data and at least one non-missing image derived phenotype value from data downloaded from UK Biobank in November 2018. ^b^The number of samples with at least one non-missing image derived phenotype value from data downloaded from UK Biobank in November 2018. ^c^Number of samples in 3 pre-defined regions of a plot of the first two genetic principal component scores, where the regions are selected to represent African, East Asian, and European ancestry (Sup. Figure 2).

| **Demographic and Clinical Characteristics** | **UKB 50k WES Participants** | **UKB 500k Participants** |
| --- | --- | --- |
| N | 49,960 | 502,543 |
| Female, n(%) | 27,243 (54.5) | 273,460 (54.4) |
| Age at assessment, years (1st-3rd Quartiles) | 58 (45-71) | 58 (45-71) |
| Body mass index, kg/m<sup>2</sup> (1st-3rd Quartiles) | 26 (21-31) | 26 (21-31) |
| Number of imaged participants (%) | 12,075 (24.1)<sup>a</sup> | 21,407 (4.3)<sup>ab</sup> |
| Number of current/past smokers, n(%) | 17,515 (35.0) | 216,482 (43.1) |
| Townsend Deprivation Index (1st-3rd Quartiles) | -2.0 (-6.1, -2.1) | -2.13 (-6.2, -1.9) |
| Inpatient ICD10 codes per patient | 5 | 5 |
| Patients with >=1 ICD10 diagnoses, n(%) | 42,066 (84.2) | 391,983 (78.0) |
| **Genetic Ancestry Assignment<sup>c</sup>** |  |  |
| African (%) | 1.49 | 1.24 |
| East Asian (%) | 0.54 | 0.51 |
| European (%) | 93.6 | 94.5 |
| **Cardiometabolic phenotypes** |  |  |
| Coronary Disease, n(%) | 3,340 (6.7) | 35,879 (7.1) |
| Heart Failure, n(%) | 300 (0.6) | 4,399 (0.8) |
| Type 2 Diabetes, n(%) | 1,541 (3.0) | 17,261 (3.4) |
| **Respiratory and immunological phenotypes** |  |  |
| Asthma, n(%) | 8,250 (16.5) | 68,149 (13.5) |
| COPD, n(%) | 741 (1.4) | 7,438 (1.4) |
| Rheumatoid Arthritis, n(%) | 710 (1.4) | 7,337 (1.4) |
| Inflammatory Bowel Disease n(%) | 543 (1.0) | 5,783 (1.1) |
| **Neurodegenerative phenotypes** |  |  |
| Alzheimer's Disease, n(%) | 13 (0.05) | 320 (0.06) |
| Parkinson's Disease, n(%) | 65 (0.13) | 1,043 (0.21) |
| Multiple Sclerosis, n(%) | 126 (0.25) | 1,352 (0.26) |
| Myasthenia Gravis, n(%) | 14 (0.02) | 217 (0.04) |
| **Oncology phenotypes** |  |  |
| Breast Cancer, n(%) | 1,667 (3.3) | 16,887 (3.3) |
| Ovarian Cancer, n(%) | 162 (0.3) | 1,777 (0.3) |
| Pancreatic Cancer, n(%) | 602 (1.2) | 4,611 (0.9) |
| Prostate Cancer, n(%) | 848 (1.6) | 8,855 (1.7) |
| Melanoma, n(%) | 598 (1.1) | 5,715 (1.1) |
| Lung Cancer, n(%) | 172 (0.3) | 2,581 (0.5) |
| Colorectal Cancer, n(%) | 368 (0.7) | 3,971 (0.8) |
| Cutaneous squamous cell carcinoma, n(%) | 1,316 (2.6) | 12,969 (2.6) |
| **Enhanced measures** |  |  |
| Hearing test, n(%) | 40,546 (81.1) | 167,011 (33.2) |
| Pulse Rate, n(%) | 40,548 (34.2) | 170,761 (33.9) |
| Visual Acuity Measured, n(%) | 39,461 (78.9) | 117,092 (23.2) |
| IOP measured (left), n(%) | 37,940 (75.9) | 111,942 (22.2) |
| Autorefracton, n(%) | 36,067 (72.1) | 105,989 (21.0) |
| Retinal OCT, n(%) | 32,748 (65.5) | 67,708 (13.4) |
| ECG at rest, n(%) | 10,829 (27.1) | 13,572 (2.1) |
| Cognitive Function, n(%) | 9,511 (19.0) | 96,362 (19.1) |
| Digestive Health, n(%) | 13,553 (28.1) | 142,310 (28.3) |
| Physical Activity Measurement, n(%) | 10,684 (21.3) | 101,117 (20.1) |

### Summary and Characterization of Coding Variation from WES

Exomes were captured using a slightly modified version of the IDT xGen Exome Research Panel v1.0. The basic design targets 39 megabases of the human genome (19,396 genes among autosomes and sex chromosomes) and was supplemented with additional probes to boost coverage at under-performing loci. In each sample and among targeted bases, coverage exceeds 20X at 94.6% of sites on average (standard deviation 2.1%). We observe 4,735,722 variants within targeted regions (Table 2). These variants include 1,229,303 synonymous (97.9% with minor allele frequency, MAF<1%), 2,498,947 non-synonymous (98.9% with MAF<1%), and 231,631 predicted LOF variants affecting at least one coding transcript (initiation codon loss, premature stop codons, splicing, and frameshifting indel variants; 99.6% with MAF<1%) (Fig. 1a). Our tally of the median number of variants per individual includes 9,619 synonymous (IQR 128), 8,781 missense (IQR 137) and 219 LOF variants (IQR 16) and is comparable to previous exome sequencing studies^12,13^; the increasing proportion of rare variants in the LOF and missense categories is consistent with purifying selection. If we restrict analysis to LOF variants that affect all ENSEMBL 85 transcripts for a gene, the number of LOF variants drops to 153,903 overall and 111 per individual (a reduction of ~33.5% and ~49.3%, respectively), consistent with previous studies. In addition to variants in targeted regions, we also capture exon adjacent variation. Including non-targeted regions, we observe 9,693,526 indel and single nucleotide variants (SNVs) after quality control, 98.5% with MAF<1%. These additional variants can be helpful aids for population genetic analyses and for applications such as phasing and IBD segment detection.

**Figure 1.**
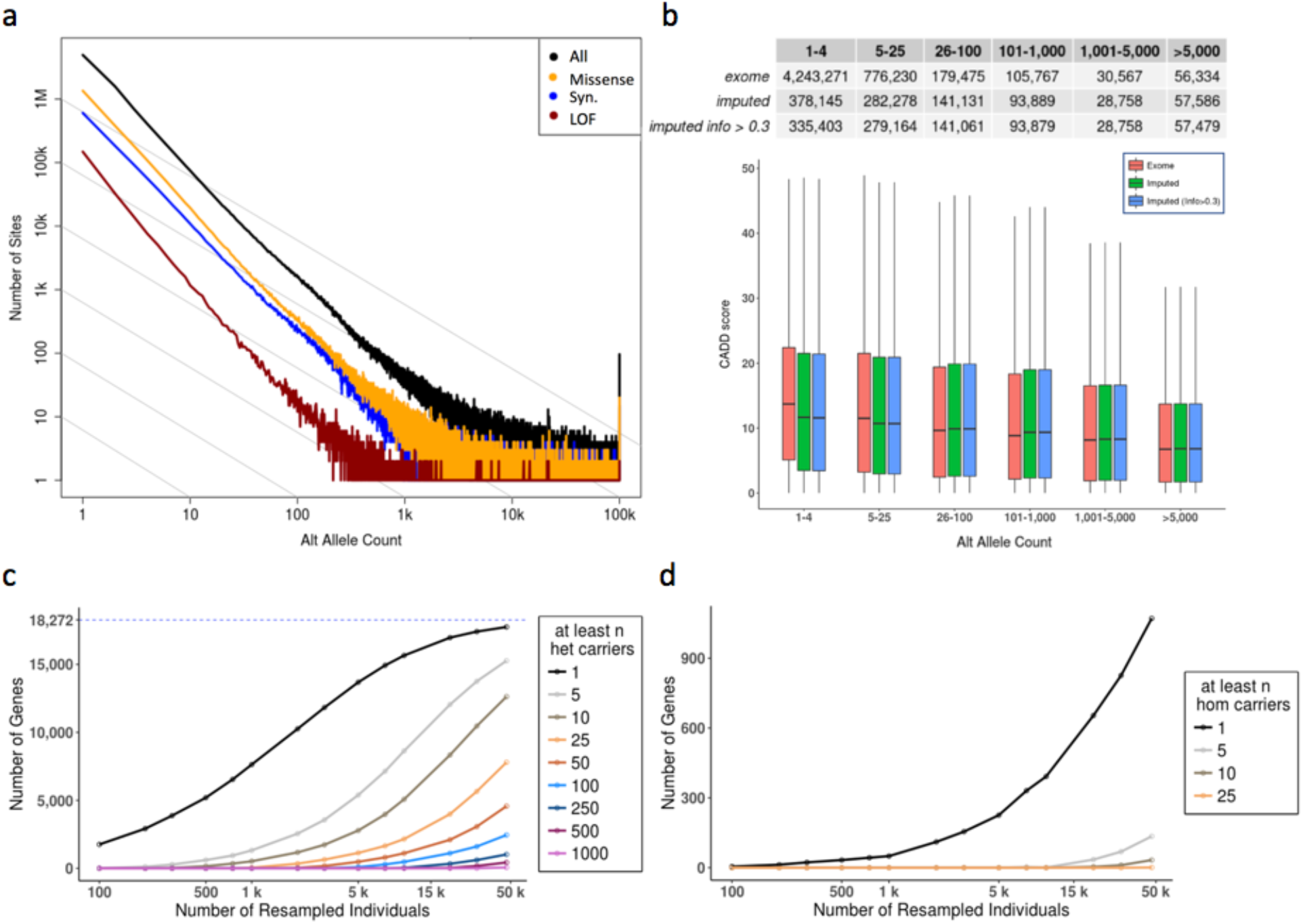
Summary statistics for variation in WES and imputed sequence a, Observed site frequency spectrum for all autosomal variants and by functional prediction in 49,960 UKB participants. **b**, Distribution of CADD scores for variant allele counts in regions consistently covered by WES (90% of individuals with >20X depth) in WES and imputed data in 49,797 UKB participants with WES and imputed sequence. **c**, The number of autosomal genes with at least 1, 5, 10, etc. heterozygous and homozygous nonreference genotypes d, LOF carriers increases with sample size. LOFs passed GL (Goldilocks) QC (see Supplemental Methods for GL QC filtering definition), genotype missingness<10%, and HWE p-value>10^−15^ 46,808 UKB participants of European ancestry with WES were down-sampled at random to the number of individuals specified on the horizontal axis. The number of genes containing at least the indicated count of LOFs MAF<1% carriers as in the legend are plotted on the vertical axis. The maximum number of autosomal genes is 18,272 in this analysis (See Supplemental Methods for gene model).

**Table 2.** Summary statistics for variants in sequenced exomes of 49,960 UKB participants. Counts of autosomal variants observed across all individuals by type/functional class for all and for MAF<1% frequency. The number of targeted bases by the exome capture design was n=38,997,831. All variants passed quality control (QC) criteria (See Supplemental Methods), individual and variant missingness <10%, and Hardy Weinberg p-value>10^−15^. Median count and interquartile range (IQR) per individual for all variants, and for MAF<1%.^1^ Counts restricted to WES targeted regions.

|  | Variants in WES, n=49,960<br>Participants |  | Median Per Participant (IQR) |  |
| --- | --- | --- | --- | --- |
|  | # Variants | # Variants<br>MAF<1% | # Variants | # Variants<br>MAF<1% |
| Total | 9,693,526 | 9,547,730 | 48,982 (627) | 1,626 (133) |
| Targeted Regions<sup>1</sup> | 4,735,722 | 4,665,684 | 24,332 (283) | 780 (63) |
| **Variant Type<sup>1</sup>** |  |  |  |  |
| SNVs | 4,520,754 | 4,453,941 | 23,529 (276) | 739 (61) |
| Indels | 214,968 | 211,743 | 803 (29) | 42 (10) |
| Multi-Allelic | 591,340 | 580,728 | 3,388 (63) | 117 (18) |
| **Functional Prediction** |  |  |  |  |
| Synonymous | 1,229,303 | 1,203,043 | 9,619 (128) | 228 (28) |
| Missense | 2,498,947 | 2,472,384 | 8,781 (137) | 380 (39) |
| LOF (any transcript) | 231,631 | 230,790 | 219 (16) | 24 (8) |
| LOF (all transcripts) | 153,903 | 153,441 | 111 (12) | 15 (6) |

### Enhancement of Coding Variation from WES Compared to Imputed Sequence

To evaluate enhancements to the UKB genetic variation resource through WES, we compared the number of coding variants observed in 49,797 individuals in whom both WES and imputed sequence were available; this is a subset of the 49,960 individuals with WES. To capture as many variants in the overlap of WES and imputed sequence as possible, only minimal filtering was applied (WES sites were filtered using genotype likelihoods, with no filtering for imputed data), thus variant counts are greater than in Table 2 and Sup. Table 1, respectively. Among all autosomal variants, we observed increases in the total number of coding variants (3,995,794 to 707,124); synonymous (1,241,804 to 267,479), missense (2,518,075 to 420,194), and LOF (235,915 to 19,451) in WES compared to imputed sequence, respectively (Sup. Table 2). This represents a >10-fold increase in the number of LOF variants identified by WES compared to those in the imputed sequence.

Amongst nearly four million coding variants observed in the exome sequence data, only 13.7% were also observed in the imputed sequence, highlighting the added value of exome sequencing for ascertainment of rare coding variation. Similarly, among LOFs observed in either dataset, 92.0% (n=223,427) were unique to WES and absent in the imputed sequence data. We observed 12,488 LOFs present in both datasets, meaning that only 5.3% of the >235,915 LOFs identified by WES were present in the imputed sequence. Since LOFs are especially informative for human genetics and medical sequencing studies, this enhancement clearly emphasizes the value of exome sequencing.

There were 6,963 LOFs seen only in the imputed sequence. These represent both variants within regions not targeted or captured in WES and errors in imputation, which are especially pronounced at low allele frequencies. Amongst the 6,963 LOFs (5,939 SNVs) observed only in the imputed sequence, we identified 1,730 LOF variants (24.8%, including 1,348 SNVs) in regions that were not targeted, 4,438 LOF variants (63.7%, including 3,804 SNVs) that fell within consistently covered regions of WES (>20x coverage in 90% of samples), and 885 LOF variants (12.7%, including 787 SNVs) in inconsistently covered exome-captured target or buffer regions. Amongst the set of 4,438 LOF variants in consistently covered regions seen solely in the imputed sequence, we selected 363 variant sample-sites from across the MAF spectrum and manually reviewed the underlying read data in the Integrative Genomics Viewer^14^ (IGV) (see Supplemental Methods). We observed that 76% of the selected sample sites had no sequencing evidence to support the imputed LOF call. Approximately 21% had some evidence for the presence of any variation (e.g. multi-nucleotide polymorphism). Only ~3% had any clear evidence of the LOF variant called in the imputed data. Sites that validated were more likely to be common than rare, as expected for imputed variants.

We also noted that amongst all the 707,124 imputed coding variants, 22.6% of them were not observed in the exome sequence data; a large portion of these will similarly suffer from poor imputation accuracy as observed in WES and imputed sequence concordance (Figure 2). As expected, common variants across functional prediction classes were more likely to be captured by both WES and imputed sequence, whereas rare variants were more likely unique to WES (Sup. Table 2). As an expected result of purifying selection, we observed that lower frequency variants were predicted to be more deleterious as measured by CADD^15^ score distributions in both datasets (Figure 1b). Interestingly, among rare variants, those identified by WES were typically classified as more deleterious (Figure 1b) – likely because rare variants that can be imputed may often be common in other populations even when rare in UKB.

**Figure 2.**
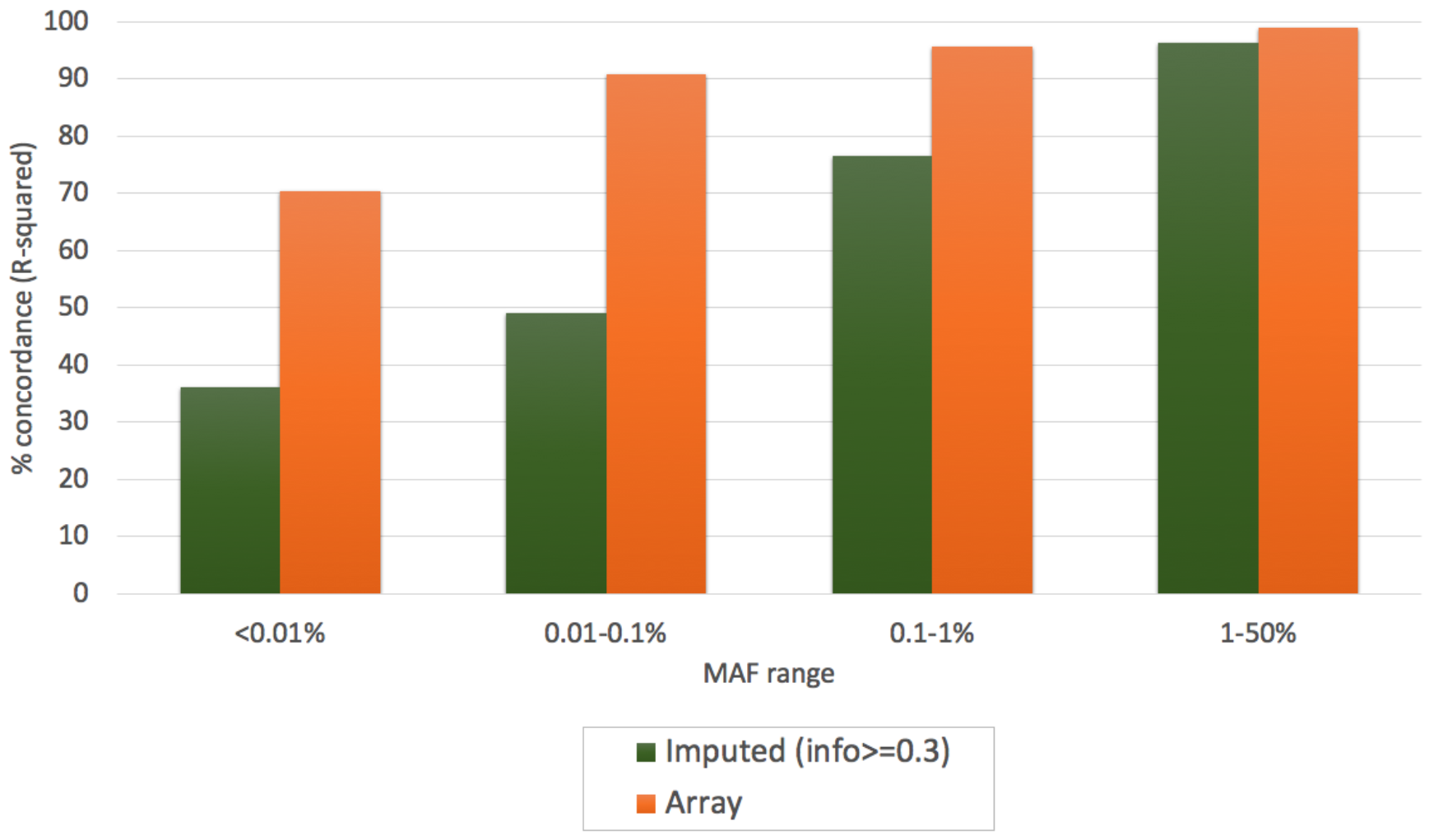
Concordance between WES, imputed sequence, and array genotypes. R-squared correlation coefficients between variants in WES and imputed sequence (green) and array genotypes and WES (orange), calculated per variant and binned by minor allele frequency in WES. n=46,912 individuals and n=75,334 variants were represented in the array-WES comparison (n=4,261, n=17,780, n=23,087 and n=30,206 variants in <0.01%, 0.01-0.1%, 0.1-1%, and 1-50% MAF bins respectively). n=46,860 individuals and n=899,455 variants were represented in the imputed-WES comparison (n=346,826, n=304,524, n=126,554, and n=121,551 variants in <0.01%, 0.01-0.1%, 0.1-1%, and 1-50% MAF bins respectively).

### Concordance of WES, Directly Genotyped Array, and Imputed Sequence

We measured concordance between WES and array genotypes in individuals with both datasets (Supplemental Methods), using the squared correlation (R^2^) of allele counts in sequencing and array genotyping. This measure facilitates interpretation of assessments of accuracy for both rare and common variation^16,17^. Using the same approach, we also measured concordance between WES and imputed sequence. As expected, concordance between genomic measures declines with decreasing MAF (Figure 2, Sup. Figures 3,4). Concordance between WES and imputed sequence ranged from 35% for MAF <0.01% to 96% for allele frequencies >1%, averaging 54% across all allele frequencies. Concordance between WES and array genotyped variants was substantially higher, ranging from 70% for MAF <0.01% to 99% for MAF >1% (Figure 2), and averaging 99% across all allele frequencies. As expected, WES performs much better in terms of concordance with array genotypes, since both directly assay the variation rather than make a computational prediction. This is particularly true in the rarest allele frequencies where the accuracy of imputation is limited using current imputation reference panels.

### Comparison of LOF Variants Ascertained Through WES and Imputed Sequence

LOF variants constitute a major class of genetic variation of great interest due to their disruption of gene function, their causal role in many Mendelian disorders, and the success of leveraging protective LOF variants to identify novel drug targets^5,6,18^. Rare LOF variation is best captured by direct sequencing approaches, such as WES, and we sought to quantify the improved yield of LOF variation from WES compared to array genotyping and imputed sequence. We compared the number of LOF variants ascertained through WES and imputed sequence among 49,797 UKB participants. Notably, not all individuals with WES have imputed sequence available. We observed a larger number of LOF variants impacting any transcript in WES vs imputed sequence, 235,915 and 19,451, respectively (See Sup. Table 2). Further, we observed a greater number of genes with ≥1 heterozygous LOF variant carrier (17,751 genes from WES, 8,763 from imputation (info >0.3) and genes with ≥1 homozygous LOF variant carrier (1,071 from WES, 789 from imputation) (Table 3). The number of genes with LOFs at different thresholds of imputation accuracy for 50k and 500k resources are included in Sup. Table 3. At equivalent sample sizes (n=46,827 European ancestry individuals with both WES and imputed sequence), WES data included a greater number of genes with LOF variants, across all carrier count thresholds. Even more striking was that WES in 46,827 individuals yielded more genes (17,751) with heterozygous LOFs than imputed sequence (info>0.3) in all 462,427 UKB participants of European ancestry (8,724 genes). Tracking the increase in the number of genes with heterozygous LOF variant carriers with the increase in the number of sequenced samples suggests that we are approaching saturation for this metric, having likely observed at least one heterozygous LOF variant carrier in most of the genes that tolerate these variants, and most genes overall (Figure 1c). In contrast, the number of genes for which homozygous LOF variants are observed still appears to increase rapidly as more samples in UKB are sequenced, suggesting that homozygous instances of LOF variants for many more genes can be identified by sequencing additional individuals (Figure 1d).

**Table 3.** Number of autosomal genes with heterozygous, homozygous LOF variants. Count of genes with at least the specified number of LOFs (MAF < 1%) impacting any transcript in European ancestry in approximately 50k (n=46,827 with WES and imputed sequence), and 462,427 individuals for 500k imputed sequence.

|  |  | # Autosomal genes containing at least N LOFs, MAF < 1% |  |  |  |  |  |
| --- | --- | --- | --- | --- | --- | --- | --- |
| Zygosity | Genomic resource | 1 | 5 | 10 | 25 | 50 | 100 |
| Het | 50k exome | 17,751 | 15,269 | 12,629 | 7,799 | 4,573 | 2,453 |
|  | 50k imputed (info > 0.3) | 8,763 | 6,999 | 5,933 | 4,297 | 2,965 | 1,911 |
|  | 500k imputed (info > 0.3) | 8,724 | 7,711 | 7,267 | 6,539 | 5,847 | 4,916 |
| Hom | 50k exome | 1,071 | 135 | 33 | 1 | 0 | 0 |
|  | 50k imputed (info > 0.3) | 789 | 74 | 7 | 0 | 0 | 0 |
|  | 500k imputed (info > 0.3) | 1,752 | 597 | 351 | 120 | 21 | 3 |

Extrapolating, we can estimate the number of genes where multiple loss of function carriers will be observed once our experiment is complete and all ~500,000 participants are sequenced. Cautiously, we currently predict that >17,000, >15,000, and >12,000 genes will have ≥10, ≥50, and ≥100 heterozygous LOF carriers in the full dataset (see Sup. Methods, Sup. Methods Fig. 1 & Table 1).

Our WES results are consistent with those of a recent large-scale survey^19^ of genetic variation in 141,456 individuals from the Genome Aggregation Database (gnomAD). When we annotate both exome variant lists with the same annotation pipelines and subset results to similar numbers of individuals and ancestry, we observe 17,751 genes with LOFs in any transcript in 46,979 European individuals in UKB with WES vs 17,946 genes with LOFs in any transcript in 56,885 Non-Finnish European (NFE) individuals in gnomAD exomes. Further subsetting to high confidence LOF variants (with LOFTEE, see Sup. Methods), we obtain 17,640 genes with LOFs in UKB and 17,856 in gnomAD (Sup. Methods Table 2).

### Survey of Medically Actionable Pathogenic Variants

To date, more than 5,000 genetic disorders have been described for which the molecular causes and associated genes have been defined. The American College of Medical Genetics (ACMG) has proposed 59 genes, ACMG SF2.0,^20,2l^ which are associated with highly penetrant disease phenotypes and for which available treatments and/or prevention guidelines can significantly reduce the morbidity and mortality in genetically susceptible individuals. Large-scale human genomic sequencing efforts coupled with EHR data provide opportunities to assess penetrance and prevalence of pathogenic (P) and likely pathogenic (LP) variants in known monogenic disease genes, as well as investigate the phenotypic effects of variants of unknown significance (VUS). Additionally, phenotypically agnostic population sampling provides opportunities to better characterize the phenotypic spectrum of these disorders and estimate the associated disease risk in the population. Furthermore, these efforts enable the implementation of precision medicine by identifying individuals carrying medically actionable pathogenic variants and providing medical care and surveillance preemptively.

We interrogated variant data for the 49,960 individuals with WES from the UKB to identify a “strict” set of reported pathogenic missense and LOF variants (that is, those with ≥2 stars in ClinVar and no conflicting interpretations) as well as a set of likely pathogenic LOF variants (that is, those in genes where truncating mutations are known to cause disease) according to the current ACMG 59 gene set^21^ (see Supplemental Methods). We identified a total of 555 such variants (316 in the reported pathogenic set, 239 in the likely pathogenic set) in 1,000 unique individuals (Table 4, Ext. Data ACMG59Variants.xlsx, 9 individuals carried variants in 2 genes). Of note, 47 of the likely pathogenic variants would qualify as previously reported pathogenic variants using a broader definition (Sup. Table 4). Variants were observed in 48 of 59 ACMG genes, with a median number of 5 variants per gene and a median 2 carriers per gene. Overall, 2.0% of the sequenced individuals carry a flagged variant in one of the ACMG59 genes. Using the same methodology in 91,514 participants from the Geisinger-Regeneron DiscovEHR study^22^ sequenced to date, we observed a slightly higher percentage of individuals, 2.76%, carrying a potentially actionable rare pathogenic variant in the ACMG59 genes (Sup. Table 5). This difference may reflect differences between a study of individuals seeking clinical care (DiscovEHR) and a population-based study not ascertained in the context of active clinical care (UKB).

**Table 4.** Medically actionable variants in ACMG59 genes in UKB participants. Of the 49,960 UKB participants with WES data, 2.0% are carriers of pathogenic (P) and likely pathogenic (LP) variants in ACMG59 v2.0 genes based on strict variant filtering criteria. LP variant counts include LOF variants passing QC criteria in ACMG59 genes that are not reported in ClinVar (≥2 star). Amongst all P+LP variants, 384 variants were observed in only one individual, 165 were observed in 2-10 individuals, and 6 were observed in >10 individuals. Percent of individuals with P or LP variants is not additive, as the 2.0% represents non-redundant carriers; 9 individuals were found to have 2 medically actionable variants.

| Category | #Variants | % of Total Known ACMG59 Variants | #Carriers | % of individuals with reportable variants |
| --- | --- | --- | --- | --- |
| Pathogenic (P) | 316 | 4.23 | 694 | 1.39 |
| Likely Pathogenic (LP) | 239 | - | 315 | 0.63 |
| P + LP | 555 | - | 1,009 | 2.0<sup>1</sup> |

Among the 48 of the reportable ACMG59 genes, variants in cancer associated genes were the most prevalent in UKB with WES; *BRCA2* (93 variants, 166 carriers), *BRCA1* (40 variants, 60 carriers), *PMS2* (21 variants, 59 carriers) and *MSH6* (35 variants, 52 carriers); while variants in *LDLR,* associated with familial hypercholesterolemia [MIM #143890], were the second most prevalent (35 variants, 68 carriers). Cardiac dysfunction disorders were also highly represented mainly by variants in *KCNQ1* (25 variants, 55 carriers), *PKP2* (22 variants, 55 carriers), and *MYBPC3* (25 variants, 50 carriers) (Figure 3a). The majority of variants were identified in genes responsible for dominant conditions, for which heterozygous carriers are at risk (994 carriers); however, we also identified 6 individuals homozygous for pathogenic variants in genes associated with autosomal recessive (familial adenomatous polyposis-2 [FAP2, MIM #608456] and mismatch repair cancer syndrome [MMRCS, MIM #276300] due to biallelic pathogenic variants in *MUTYH* and *PMS2,* respectively) or hemizygous for X-linked conditions (Fabry disease [MIM #301500] due to pathogenic variants in *GLA).* Indeed, a male hemizygous for the c.335G>A; p.Arg112His pathogenic variant in *GLA* has diagnoses of angina pectoris, atrial fibrillation, chest pain, and chronic ischemic heart disease. Similarly, one individual homozygous for a pathogenic missense variant (c.1145G>A; p.Gly382Asp) in *MUTYH* has a history of benign neoplasm of colon, diverticular disease of the intestine, colonic polyps, and intestinal obstruction. These examples illustrate how the extensive health data available for UK Biobank participants provide a valuable resource to assess variant pathogenicity and disease risk at the population level, and the potential to model outcomes for individuals harboring pathogenic variants.

**Figure 3.**
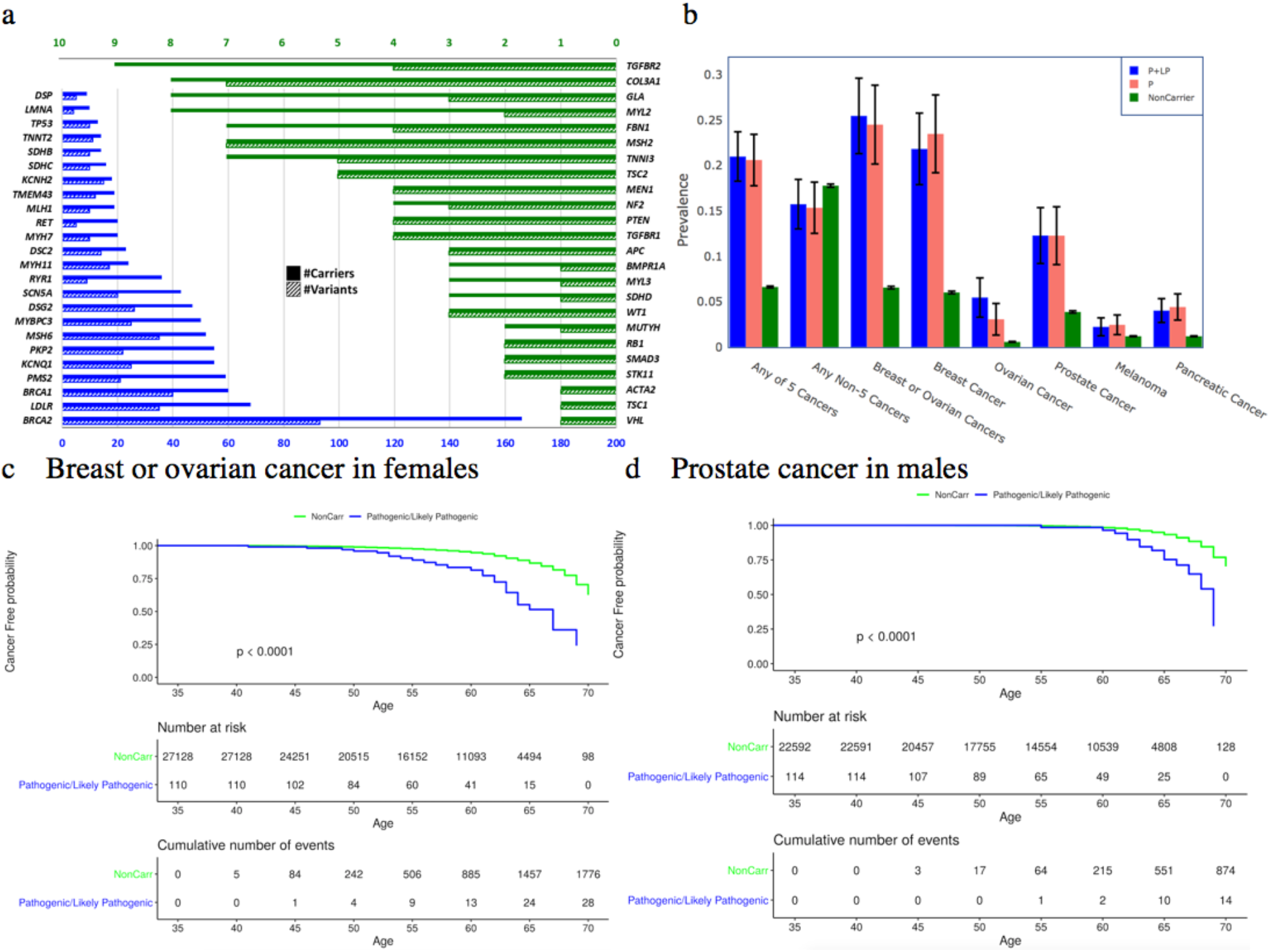
Summary of observed actionable ACMG59 variants, and pathogenicity of *BRCA1/2* variants. a, Counts of variants and carriers in 48 ACMG59 genes with pathogenic (P) or likely pathogenic (LP) variants. **b**, Prevalence of cancers in carriers of P, P or LP, and no P or LP variants in *BRCA1* or *BRCA2.* Five major cancers related to *BRCA1/2* risk include breast in females, ovarian, prostate, melanoma, and pancreatic cancers. Cases are aggregated from Cancer Registry, HES, and Self Report. **c**, Cumulative proportion of female participants free of breast and ovarian cancer, and **d**, male participants free of prostate cancer with P or LP variants in *BRCA1/2* compared to non-carriers by age at interview.

As an example, we evaluated the risk of cancer in individuals carrying pathogenic variants in *BRCA1* or *BRCA2* (Figure 3b) to compare across five previously implicated *BRCA1/2* associated cancers^23^ as well as to explore whether risk was conferred for other cancers. While *BRCA1* and *BRCA2* have differences in mechanism and risk among cancer subtypes, due to low sample sizes, we analyzed summed counts in 114 males and 110 females with *BRCA1/2* reported pathogenic and likely pathogenic variants. We found the prevalence of cancers to be elevated in carriers of pathogenic variants in *BRCA1/2* for the 5 cancers (breast in females, ovarian, prostate, melanoma, and pancreatic) previously associated with *BRCA1* or *BRCA2* (cancer status was derived from self-report, HES, and cancer prevalence for any of the 5 cancers was 21.0% in carriers vs 6.6% in non-carriers, OR = 3.75 (95%CI 2.71,5.18), p-value = 2×10^−12^, (3337 cases, 46599 controls), and there was no significant difference between carriers and non-carriers when all other cancers, excluding these 5, were combined; 15.7% in carriers vs 17.7% in non-carriers, OR = 0.86 (95%CI 0.58,1.29), p-value = 0.55, (8300 cases, 38424 controls). The most prevalent cancers in this group were C443 and C449 unspecified malignant neoplasms of skin, and D069 carcinoma in situ of cervix. Increased risk was observed in these *BRCA1/2* variant carriers for each of the five previously associated cancers analyzed individually (ovarian in females OR=9.97 (95%CI 4.32,23.06), p=5.2×10^−5^, (162 cases, 27,076 controls); breast in females OR=4.37 (95%CI 2.77,6.88), p=3.6×10^−8^, (1,654 cases, 25,584 controls); prostate in males OR=3.48 (95%CI 1.98, 6.11), p=1.5×10^−4^, (888 cases, 21,818 controls); melanoma OR=1.9 (95%CI 0.78,4.62), p=0.20, (599 cases, 49,345 controls); and pancreatic cancer OR 3.47 (95%CI 1.77,6.79), p=1.7×10^−3^, (602 cases, 49,342) controls; (p values by Fisher’s exact test). These differences in overall cancer risk also manifest as clearly different disease onset and rates of cancer-free survival between carriers and non-carriers (see Figure 3c for breast and ovarian cancer in females; Figure 3d for prostate cancer in males). Comparing the cumulative proportion of female participants free of breast and ovarian cancer, we estimate a hazard ratio of 4.3 (95%CI 2.96,6.24, p<0.0001). Comparing the cumulative proportion of male participants free of prostate cancer, the hazard ratio was 3.68 (95%CI 2.17, 6.24, p<0.0001). With the UKB resource, cancer risk can be more deeply explored across broader sets of variants as well as with larger exome sequence datasets and continued accrual of incident cancers. Our results corroborate those of another recent population-based application of WES linked to health records to evaluate cancer risk in individuals with pathogenic variants in *BRCA1/2*^10^, demonstrating the value of WES to identify high-penetrance rare alleles associated with clinical phenotypes; such efforts can be applied across other genetic disorders, enabling the implementation of precision medicine at the population level.

### Phenotypic Associations with LOF Variation

The combination of WES, allowing comprehensive capture of LOF variants, with rich health information allows for broad investigation of the phenotypic consequences of human genetic variation. We conducted burden tests for rare (MAF < 1%) LOF variants in autosomal genes with >3 LOF variant carriers and in 1,741 traits (1,073 discrete traits with at least 50 cases defined by HES and self-report data, 668 quantitative, anthropometric, and blood traits) in 46,979 individuals of European ancestry. For each gene-trait association, we also evaluated signal for the single variants included in the burden test. We identified 25 unique gene burden-trait associations with p < 10^−7^; among these 21 were more significant than any single LOF variant included in the burden test. The results include several well-established associations (Table 5). For example, we observe that carriers of *MLH1* LOFs, associated with Muir-Torre and Lynch syndromes^24^ [MIM #158320, #609310], were at increased risk of malignant neoplasms of the digestive organs (OR=84, P=3.5×10^−11^). Carriers of *PKD1* LOFs, the major cause of autosomal dominant polycystic kidney disease^25^ [MIM #173900], were at increased risk of chronic kidney disease (OR=91, P=2.9×10^−10^). Carriers of *TTN* LOFs are at increased risk for cardiomyopathy (OR=11.9, P=1.4×10^−8^), consistent with prior reports^26^. In addition to Mendelian disorders, other findings with strong support in the literature include *HBB* with red blood cell phenotypes^27^, *IL33* with eosinophils (driven by rs146597587)^28^, *KALRN* with platelet volume (driven by rs56407180)^29^, *TUBB1* with multiple platelet phenotypes^30^, and *CALR* with hematopoietic neoplasms^31^. In some cases, we see patterns of association with traits that may be secondary to known phenotypic associations. For example, *ASXL1* and *CHEK2* are genes involved in myeloproliferative disorders^32^ and cancer^33^, respectively, which may explain the observed associations with hematologic traits (which may be secondary to myelodysplastic disease or chemotherapy). Many other known phenotypic associations are supported by the data at more modest significance thresholds (Sup. table 6). These include, for example, associations between LOF variants in *LDLR* with coronary artery disease, *GP1BB* with platelet count, *PALB2* with cancer, and *BRCA2* with cancer risk.

**Table 5.**
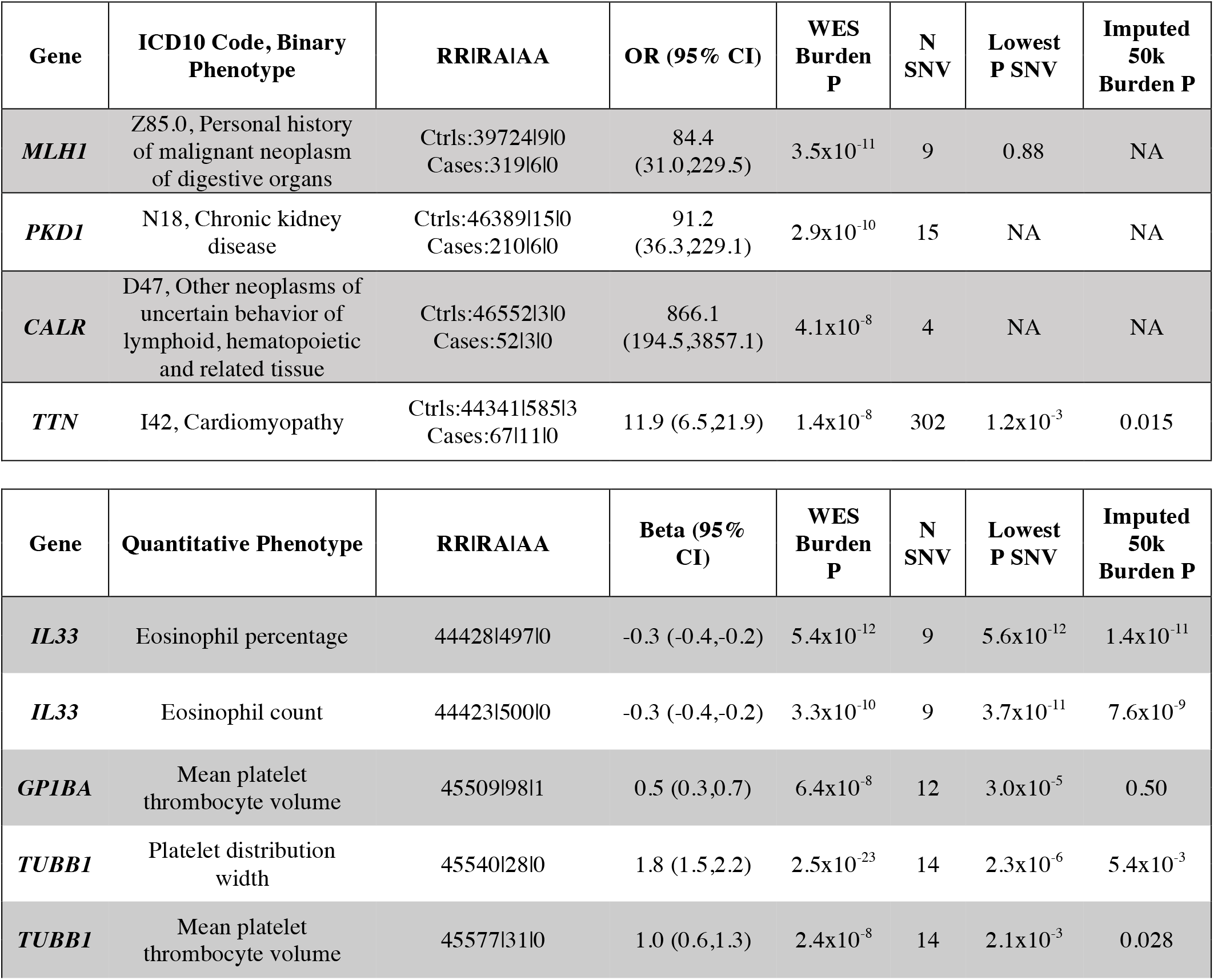
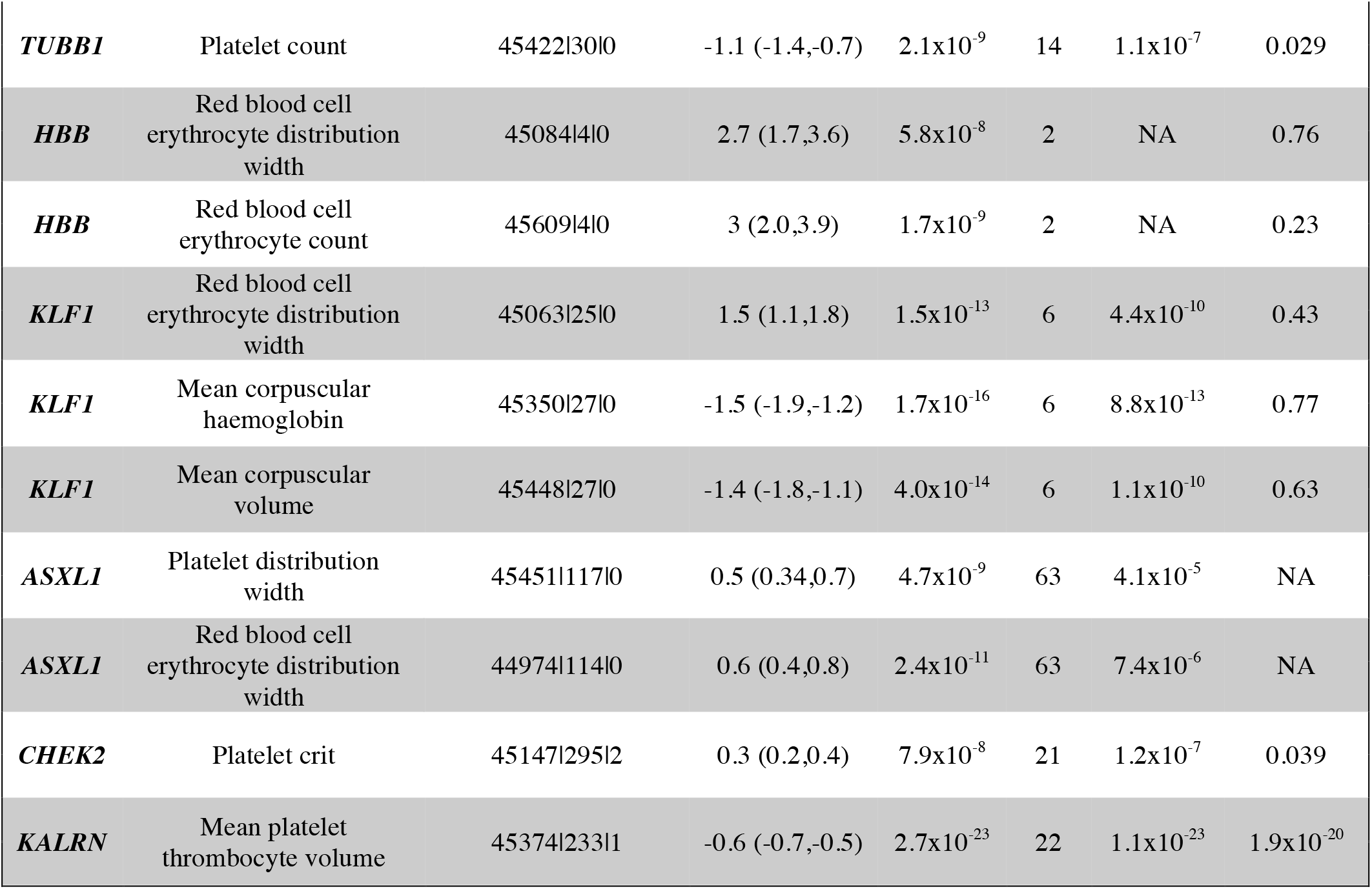
LOF gene burden results with previously known genetic associations. LOF gene burden association with available clinical and continuous traits in 46,979 UKB participants of European ancestry with WES.

### LOF Associations and Novel Gene Discovery

Our LOF gene burden association analysis identified five gene-trait LOF associations with p<10^−7^ that have not previously been reported (Table 6). We identified a novel association between *PIEZO1* LOFs (cumulative allele frequency = 0.2%) and increased risk for varicose veins (OR=4.8, P=2.7×10^−8^). This finding is driven by a burden of rare LOF variants, with the most significant *PIEZO1* single variant LOF association achieving a p-value of 2.3×10^−3^. ‘Leave-one-out’ (LOO) analyses indicated no single variant accounted for the entire signal and step-wise regression analyses indicated that 11 separate variants (5 of which had minor allele count (MAC)>1) were contributing to the overall burden signal (Sup. Table 8 and Ext. Data SingleVariantLOFs.xlsx). We replicated this finding for varicose veins (1,572 cases, 75,704 controls) in WES data from the DiscovEHR study (OR= 3.8, p= 1.5×10^−6^) (Sup. Table 10). This region had previously been implicated by common non-coding variants with small effects^34^ (rs2911463, OR = 0.996; Supplemental Table 9). *PIEZO1* encodes a 36 transmembrane domain cation channel that is highly expressed in the endothelium and plays a critical role in the development and adult physiology of the vascular system, where it translates shear stress into electrical signals^35^. This previous report of the rs2911463 variant mapped the association to *PIEZO1* through evidence of gene function and analysis using DEPICT, but did not find strong eQTL evidence that would clarify mechanism to modulate the target therapeutically^34^. Rare missense variants have been reported in families segregating autosomal dominant dehydrated hereditary stomatocytosis (DHS, MIM #194380) characterized by hemolytic anemia with primary erythrocyte dehydration due to decreased osmotic fragility. Whereas biallelic loss of function variants in *PIEZO1* have been reported in families with lymphatic malformation syndrome [MIM #616843], a rare autosomal recessive disorder characterized by generalized lymphedema, intestinal and/or pulmonary lymphangiectasia and pleural and/or pericardial effusions. Interestingly, some of the reported patients presented with varicose veins and deep vein thrombosis^36^. Our data provide compelling support for *PIEZO1* as the causal gene in this locus, but also clarifies direction of effect in that loss of gene function in heterozygous carriers leads to increased risk of developing varicose veins.

**Table 6.** Novel LOF gene burden associations. LOF gene burden association with available clinical and continuous traits in 46,979 UKB participants of European ancestry with WES. ^l^No *COL6A1* LOFs were observed in imputed sequence.

| Gene | ICD10 Code,<br>Binary Phenotype | RR\|RA\|AA | OR<br>(95% CI) | WES<br>Burden P | N<br>SNV | Lowest<br>P SNV | Imputed<br>50k<br>Burden P |
| --- | --- | --- | --- | --- | --- | --- | --- |
| *PIEZO1* | I83.9,<br>Asymptomatic<br>varicose veins of<br>lower extremities | Ctrls:43285\|142\|0<br>Cases:1267\|20\|0 | 4.8<br>(3.1,7.8) | $2.7 \times 10^{-8}$ | 65 | $2.3 \times 10^{-3}$ | 0.083 |
| Gene | Quantitative<br>Phenotype | RR\|RA\|AA | Beta<br>(95% CI) | WES<br>Burden P | N<br>SNV | Lowest<br>P SNV | Imputed<br>50k<br>Burden P |
| *MEPE* | Heel bone mineral<br>density | 42898\|150\|0 | -0.5<br>(-0.6,-0.3) | $1.4 \times 10^{-8}$ | 16 | $6.2 \times 10^{-5}$ | 0.033 |
| *COL6A1* | Corneal resistance<br>factor mean | 35621\|17\|0 | -1.5<br>(-2.0,-1.0) | $3.6 \times 10^{-10}$ | 11 | $1.5 \times 10^{-4}$ | NA<sup>1</sup> |
| ***COL6A1*** | Corneal hysteresis mean | 35620\|17\|0 | -1.3<br>(-1.8,-0.9) | 2.1x10<sup>-8</sup> | 11 | 4.6x10<sup>-4</sup> | NA<sup>1</sup> |
| ***IQGAP2*** | Mean platelet thrombocyte volume | 45443\|165\|0 | 0.7<br>(0.5,0.8) | 1.1x10<sup>-19</sup> | 53 | 9.3x10<sup>-8</sup> | 0.10 |
| ***GMPR*** | Mean corpuscular haemoglobin | 45195\|182\|0 | 0.4<br>(0.3,0.6) | 1.1x10<sup>-8</sup> | 13 | 6.0x10<sup>-8</sup> | 6.2x10<sup>-6</sup> |

We also identified a novel LOF burden association with *MEPE* (cumulative MAF 0.18%) and decreased bone density (as measured by heel bone mineral density, −0.50 standard deviations (SD), p-value = 1.4×10^−8^). LOO analyses (Sup. Table 11) suggested that the aggregate signal is driven by multiple variants, one of which could be imputed (rs753138805, encoding a frameshift predicted to result in early protein truncation). In analysis of all 500k UKB participants, rs753138805 was significantly associated with decreased BMD (−0.4 SD, P=8.1×10^−19^) and showed a trend for increased risk of osteoporosis (N=3,495 cases, OR=1.9, P=0.10, Sup. Table 16). These findings are corroborated by a query of the HUNT study^37^, where rs753138805 was associated with decreased BMD (−0.5 SD, P=2.1×10^−18^) and increased risk of any fracture (N=24,155 cases, P=1.6×10^−5^, OR=1.4) (Sup. Table 16). In DiscovEHR, we observed a directionally consistent, but nonsignificant, association for *MEPE* LOFs with femoral neck BMD T-score (B = −0.19, P = 0.22). The *MEPE* locus was previously implicated in GWAS^38^, with six independent signals with modest effects on bone mineral density. While two of the previously reported non-coding variants are in moderate (r^2^=0.5) or high (r^2^=0.78) linkage disequilibrium (LD) with two variants contributing to the burden test (Sup. Table 12), the burden association is only partially attenuated in conditional analyses (p~2×10^−4^ with all 6 variants together). *MEPE* encodes a secreted calcium-binding phosphoprotein with a key role in osteocyte differentiation and bone homeostasis^39,40^. Studies in *Mepe^-/-^* knockout mice *(Mepe^m1Tbrw^)* have yielded inconsistent results, with two groups reporting an increase in BMD^40,41^ and another reporting no change^42^.

In another novel signal, *COL6A1* LOFs (cumulative allele frequency = 0.03%) are associated with a 2.7 mmHg decrease in corneal resistance factor (CRF) (−1.5 SD, p-value = 3.6×10^−10^) and corneal hysteresis (CH) (−1.3 SD, p-value=2.1×10^−8^), which are measures of corneal biomechanics^43^. *COL6A1* encodes a component of collagen type VI microfibrils, which play important roles in maintaining structure and function of the extracellular matrix, and which are major components of the human cornea^44^. This locus has previously been implicated in ocular traits; rs73157695 and nearby common variants have been associated with myopia^45^ (OR = 0.94; p = ~10^−13^) and intraocular pressure^46^. LOO analyses indicate multiple variants are driving the association with both corneal traits (Sup. Tables 13, 14) and that these are not in strong LD with previously reported variants (Sup. Table 15). COL6A1 protein levels were reduced in eyes from patients with keratoconus^47^, and individuals with keratoconus and other corneal diseases such as Fuchs’ corneal dystrophy have reduced CH and CRF^48^. Measures of CH and CRF were not available in DiscovEHR for replication analyses.

The remaining novel LOF associations in *IQGAP2* and *GMPR* (driven by rs147049568) are for hematologic traits in which variants in/near each gene have previously been implicated by GWAS^29,49^. Our results for these two genes, each of which replicated in DiscovEHR (Sup. Table 7), provide additional evidence for causal roles for these genes and establishes direction of effect with respect to gene function on hematologic traits. In equivalent sample sizes, LOF burden results from imputed sequence would not have uncovered the novel LOF associations at p < 10^−7^ as identified by WES (Table 6). Further, for 19 of 25 LOF burden results described herein, gene burden results from WES in 50k were more significant than burden results from imputed sequence in all 500k UKB participants (Sup. table 10), demonstrating the value of rare variants captured by WES to power these associations.

## DISCUSSION

Integration of large scale genomic and precision medicine initiatives offer the potential to revolutionize medicine and healthcare. Such initiatives provide a foundation of knowledge linking genomic and molecular data to health-related data at population scale, allowing for the ability to more completely and systematically study genetic variation and its functional consequences on health and disease. Here, we describe the initial tranche of large-scale exome sequencing of 49,960 UK Biobank participants, which to our knowledge is currently the largest open access resource of exome sequence data linked to health records and extensive longitudinal study measures. These data greatly extend the current genetic resource, particularly in ascertainment of rare coding variation, which we demonstrate has utility in resolving variant to gene links and directionality of gene to phenotype associations.

After quality control, we observed nearly four million single nucleotide and indel coding variants. Only approximately 14% of coding variants identified by WES were observed in the imputed sequence of 49,797 participants with both WES and imputed sequence, highlighting the added value of exome sequencing. This enrichment was even more pronounced with LOF variation where WES identified >230,000 LOF variants and only approximately 5% of these were present in the imputed sequence. Further, 22.6% of the coding variants in the imputed sequence were not observed in the exome sequence data which may represent a large proportion of rare variants that have poor imputation accuracy, as observed in our concordance and visual validation analyses. A small proportion of these variants, seen only in the imputed sequence, also represents variants not in the regions targeted by exome capture design and sequencing, a limitation of the targeted capture approach. Increasing numbers of individuals and ancestral diversity in imputation reference panels are expected to improve imputation accuracy for rare variants.

As with previous studies of this size^22^, we observed a large number of LOF variants, including at least one rare heterozygous LOF variant carrier in >97% of autosomal genes (compared to >36% of autosomal genes in the imputed sequence for the same participants). It is important to note that our LOF annotation strategy is geared towards increasing sensitivity for identification of LOF variants and novel downstream association discovery. While the number of genes with heterozygous instances of LOF variants is approaching saturation at this sample size, exome sequencing of the entirety of the UKB resource will dramatically increase the number of LOF carriers and the ability to detect phenotypic associations. We also observe 1,071 autosomal genes with homozygous instances of rare LOF variants, and this number of genes will also increase with continued sequencing of all UKB participants; however, studies in populations with a high degree of parental relatedness^50,51^ will provide yet more genes with homozygous LOFs and complement efforts such as UKB. LOF variation is an extremely important class of variation for identifying drivers of high genetic risk, novel disease genes, and therapeutic targets. Very large samples sizes are needed to detect novel LOF associations given their collective rare allele frequencies. The exome sequence data provides a substantial enhancement to the number of LOF variants identified and power for detecting novel associations, which will only improve with continued sequencing of all UKB participants.

We illustrate the unique value of this expanded exome sequence resource in the UKB to assess pathogenic and likely pathogenic variants in an unascertained large-scale population-based study with longitudinal follow up. We conducted a survey of pathogenic and likely pathogenic variants in the medically actionable ACMG59 genes. Using stringent variant filtering criteria, we arrived at an estimated prevalence of 2% of individuals in this study population having a clinically actionable finding. This resource allows us to characterize disease risk profiles for individuals who carry pathogenic and likely pathogenic variants in medically relevant disease genes, including cancer susceptibility genes, such as *BRCA1* and *BRCA2.* We observed that pathogenic and likely pathogenic variant carriers had 3.75-fold greater odds of any of the 5 cancers previously associated with *BRCA1/2*; prevalence for any of the 5 cancers was 21.0% in carriers vs 6.6% in non-carriers. We further explored whether these variants conferred risk to any other cancers and did not observe any such associations. This resource will be valuable for assessment of variant pathogenicity, particularly for variants of unknown significance and novel variants, and in exploring the full spectrum of disease risk and phenotypic expression. One limitation of the resource for such purposes is limited ancestral diversity. This and other similar studies also highlight the value and potential to apply large scale sequencing at the population scale to identify a meaningful proportion of individuals who are at high risk of diseases where effective interventions are available that can significantly reduce the morbidity and mortality of genetically susceptible individuals; such precision medicine approaches could substantially reduce the burden of many diseases.

We conducted gene burden association testing for LOF variants across all genes and encompassing greater than 1,700 binary and quantitative traits. In addition to replication of numerous positive controls, we also identified a handful of significant novel LOF associations highlighting novel biology and genetics of large effect on disease traits of interest; this included *PIEZO1* for varicose veins, *MEPE* for bone density, and *COL6A1* for corneal thickness, amongst others. We identified a novel burden association in *PIEZO1*, a mechanosensing ion channel present in endothelial cells in vascular walls, that confers a nearly five-fold increased odds of varicose veins in heterozygous LOF carriers. We also identified a novel LOF burden association in *MEPE* with decreased BMD and an approximately 2-fold increased odds of osteoporosis and 1.5-fold increased risk of fractures. Overall, through WES and gene burden tests of association for LOF variants, we identified 25 unique gene-trait associations exceeding a p<10^−7^ of which 21 were substantially more significant than any single LOF variant included in the burden test, highlighting the value of WES and the ability to detect novel associations driven by rare coding variation. While these regions had previously been identified in genome-wide association studies of >10x the sample size, a key strength of the current approach is compelling identification of likely causal genes and the direction of effect: two key pieces of information required for translation towards novel therapeutics. This survey of rare LOF associations was limited by sample size for most binary traits but was well powered for many quantitative traits. While surveys of LOF variation in the entire UKB study using array and imputed sequence have identified LOF associations in previous reports^52,53^, WES identifies novel associations, unique to exome sequence and detected in only approximately one tenth of the sample size; this highlights the considerable power of exome sequencing for LOF and rare variant association discovery and the further promise of novel biological insights through sequencing all participants in the UKB resource.

Efforts are underway to sequence the exomes of all 500,000 UKB participants; these efforts will greatly expand the total amount of rare coding variation ascertained, including the number of heterozygous LOF instances that can now be observed in nearly all genes and the number of genes for which naturally occurring homozygous knockouts can be observed. Coupled with rich laboratory, biomarker, health record, imaging, and other health related data continually added to the UKB resource, exome sequencing will enhance the power for discovery and will continue to yield many important findings and insights. The WES data is available to approved researchers through similar access protocols as existing UK Biobank data (see URLs).

## Supporting information

ACMG59Variants.xlsx

HESinWESvs500k_V1.xlsx

SingletonpLOFVizVal_V1.xlsx

SingleVariantLOFs_V1.xlsx

TraitsLists.xlsx

## ACKNOWLEDGEMENTS

The authors thank the UKBiobank participants and researchers for creating an open scientific resource for the research community; the MyCode Community Health Initiative participants for taking part in the DiscovEHR collaboration; and the participants of the Nord-Trøndelag Health Study (HUNT), a collaboration between the HUNT Research Centre (Faculty of Medicine, Norwegian University of Science and Technology NTNU), the Nord-Trøndelag County Council, the Central Norway Health Authority and the Norwegian Institute of Public Health. This work was referring UKB application 26041.

## URLs

UK Biobank website and data access http://ukbiobank.ac.uk/

Reprints and permissions information is available at www.nature.com/reprints

## AUTHOR INFORMATION

### Contributions

C.V.H., M.R.N., J.W., J.G.R., J.M., J.D.O., R.A.S., L.Y-A, G.A., A.B., directed and designed research; J.B., B.Y., D.L., A.H.L., A.M., C.G-J., C.O., C.V.H., I.T., J.M. contributed to statistical analyses; C.G.-J., W.C., S.B., B.Y., S.K., J.S., A.L.B., C.V.H. contributed to the medically actionable variants survey and cancers analysis, N.B., J.X.H., B.Y., A.Y., S.K., A.H., S.B., M.C., contributed to the preparation of genetic and phenotype data; E.M., L.B., A.L., L.H., J.P., W.J.S., J.G.R., J.D.O. contributed to exome sequencing and variant calling; I.S., C.J.W., K.H., J.B.L., D.J.C., D.H.L. contributed data or results for replication; C.V.H., J.B., I.T., L.Y-A., C.G.J., C.O., S.B., N.B., R.A.S., J.M., J.R., G.A., A.B. co-wrote the manuscript. All authors reviewed the manuscript.

### Competing interests

C.V.H., J.D.B., B.Y., C.G.J., S.K., D.L., N.B., A.H.L., C.O., A.M., J.S., C.S., A.H., E.M., L.B., A.L., J.P., L.H., A.L.B., A.Y., K.P., M.J., W.J.S., G.D.Y., A.E., G.C., A.R.S., S.B., M.C., J.G.R., J.M., J.D.O., G.A., A.B. are current or former employees and/or stockholders of Regeneron Genetics Center or Regeneron Pharmaceuticals.

I.T., J.X.H., A.P., L.C., M.R.N., J.W., R.A.S., L.Y-A are current or former employees and/or stockholders of GlaxoSmithKline.

L.C. is a current employee of BioMarin.

No other authors declare a competing interest.

### Regeneron Genetics Center Banner Author List and Contribution Statements

All authors/contributors are listed in alphabetical order.

#### RGC Management and Leadership T eam

Goncalo Abecasis, Ph.D., Aris Baras, M.D., Michael Cantor, M.D., Giovanni Coppola, M.D., Aris Economides, Ph.D., John D. Overton, Ph.D., Jeffrey G. Reid, Ph.D., Alan Shuldiner, M.D.

Contribution: All authors contributed to securing funding, study design and oversight, and review and interpretation of data and results. All authors reviewed and contributed to the final version of the manuscript.

#### Sequencing and Lab Operations

Christina Beechert, Caitlin Forsythe, M.S., Erin D. Fuller, Zhenhua Gu, M.S., Michael Lattari, Alexander Lopez, M.S., John D. Overton, Ph.D., Thomas D. Schleicher, M.S., Maria Sotiropoulos Padilla, M.S., Karina Toledo, Louis Widom, Sarah E. Wolf, M.S., Manasi Pradhan, M.S., Kia Manoochehri, Ricardo H. Ulloa.

Contribution: C.B., C.F., K.T., A.L., and J.D.O. performed and are responsible for sample genotyping. C.B, C.F., E.D.F., M.L., M.S.P., K.T., L.W., S.E.W., A.L., and J.D.O. performed and are responsible for exome sequencing. T.D.S., Z.G., A.L., and J.D.O. conceived and are responsible for laboratory automation. M.P., K.M., R.U., and J.D.O are responsible for sample tracking and the library information management system.

#### Genome Informatics

Xiaodong Bai, Ph.D., Suganthi Balasubramanian, Ph.D., Leland Barnard, Ph.D., Andrew Blumenfeld, Yating Chai, Ph.D., Gisu Eom, Lukas Habegger, Ph.D., Young Hahn, Alicia Hawes, B.S., Shareef Khalid, Jeffrey G. Reid, Ph.D., Evan K. Maxwell, Ph.D., John Penn, M.S., Jeffrey C. Staples, Ph.D., Ashish Yadav, M.S.

Contribution: X.B., A.H., Y.C., J.P., and J.G.R. performed and are responsible for analysis needed to produce exome and genotype data. G.E., Y.H., and J.G.R. provided compute infrastructure development and operational support. S.K., S.B., and J.G.R. provide variant and gene annotations and their functional interpretation of variants. E.M., L.B., J.S., A.B., A.Y., L.H., J.G.R. conceived and are responsible for creating, developing, and deploying analysis platforms and computational methods for analyzing genomic data.

#### Clinical Informatics

Nilanjana Banerjee, Ph.D., Michael Cantor, M.D.

Contribution: All authors contributed to the development and validation of clinical phenotypes used to identify study participants and (when applicable) controls.

#### Analytical Genomics and Data Science

Goncalo Abecasis, Ph.D., Amy Damask, Ph.D., Manuel Allen Revez Ferreira, Ph.D., Lauren Gurski, Alexander Li, Ph.D., Nan Lin, Ph.D., Daren Liu, Jonathan Marchini Ph.D., Anthony Marcketta, Shane McCarthy, Ph.D., Colm O’Dushlaine, Ph.D., Charles Paulding, Ph.D., Claudia Schurmann, Ph.D., Dylan Sun, Cristopher Van Hout, Ph.D., Bin Ye

Contribution: Development of statistical analysis plans. QC of genotype and phenotype files and generation of analysis ready datasets. Development of statistical genetics pipelines and tools and use thereof in generation of the association results. QC, review and interpretation of result. Generation and formatting of results for manuscript figures. Contributions to the final version of the manuscript.

#### Therapeutic Area Genetics

Jan Freudenberg, M.D., Nehal Gosalia, Ph.D., Claudia Gonzaga-Jauregui, Ph.D., Julie Horowitz, Ph.D., Kavita Praveen, Ph.D.

Contribution: Development of study design and analysis plans. Development and QC of phenotype definitions. QC, review, and interpretation of association results. Contributions to the final version of the manuscript.

#### Planning, Strategy, and Operations

Paloma M. Guzzardo, Ph.D., Marcus B. Jones, Ph.D., Lyndon J. Mitnaul, Ph.D.

Contribution: All authors contributed to the management and coordination of all research activities, planning and execution. All authors managed the review of data and results for the manuscript. All authors contributed to the review process for the final version of the manuscript.

