## Supplemental Figures and Tables for "Whole exome sequencing and characterization of coding variation in 49,960 individuals in the UK Biobank"

Supplemental Tables and Figures

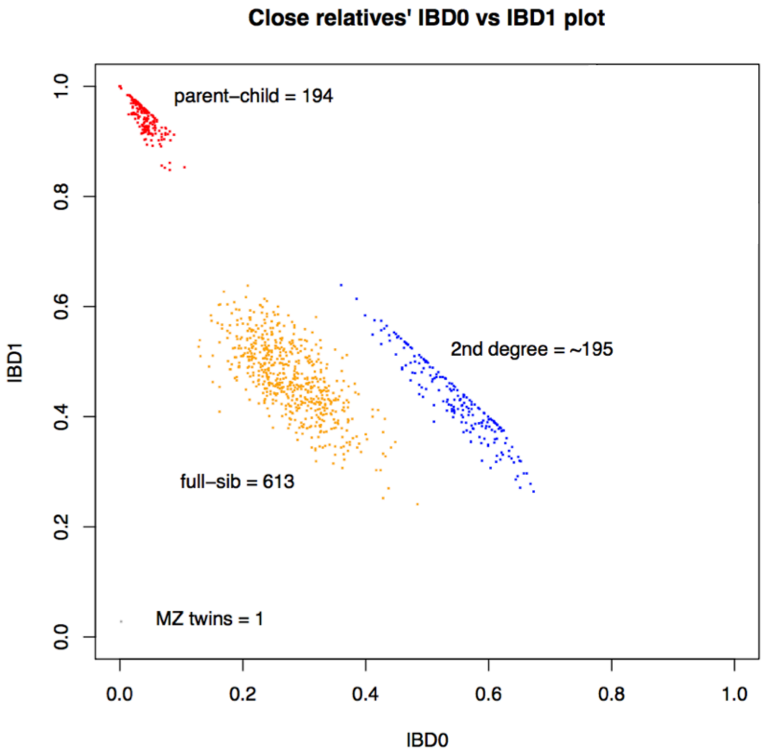


**Sup. Figure 1 | Distribution of IBD sharing for pairs of individuals in UKB 50k WES.** Estimated proportion of WES genotypes with no alleles identical by descent (IBD) vs. 1 allele IBD amongst all pairs of UKB 50k exome participants. Note that 2nd relationship pairs (blue) are included only if they are part of a relationship network containing a parent-child (red) full-sib (orange) or monozygotic twin (grey) relationship pair. We calculated IBD proportions in PLINK[1], restricting the analysis to common single nucleotide polymorphisms with minor allele frequency >10%, genotype missingness <5%, and a Hardy-Weinberg Equilibrium p-value > 0.00001.

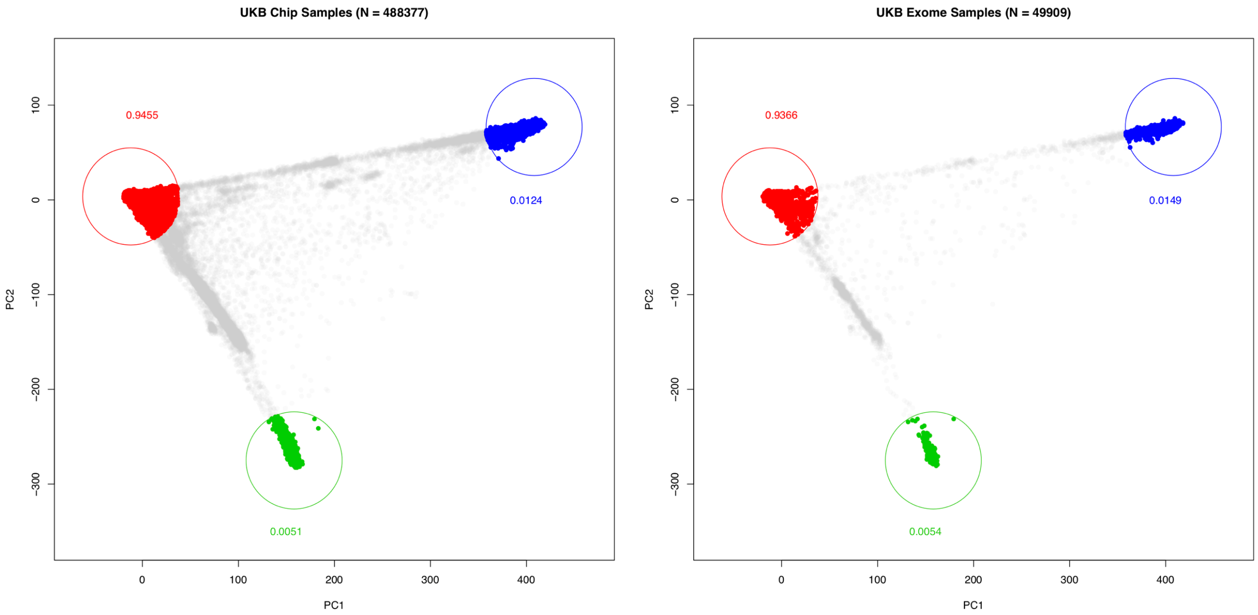


**<u>Sup. Figure 2 | Continental ancestry in UK Biobank 500k and 50k.</u>** Principal component 1 and 2 for n=488,377 individuals available from the UK Biobank Data Showcase. Three pre-defined regions of a plot of represent African (blue), East Asian (green), and European (red) ancestry.

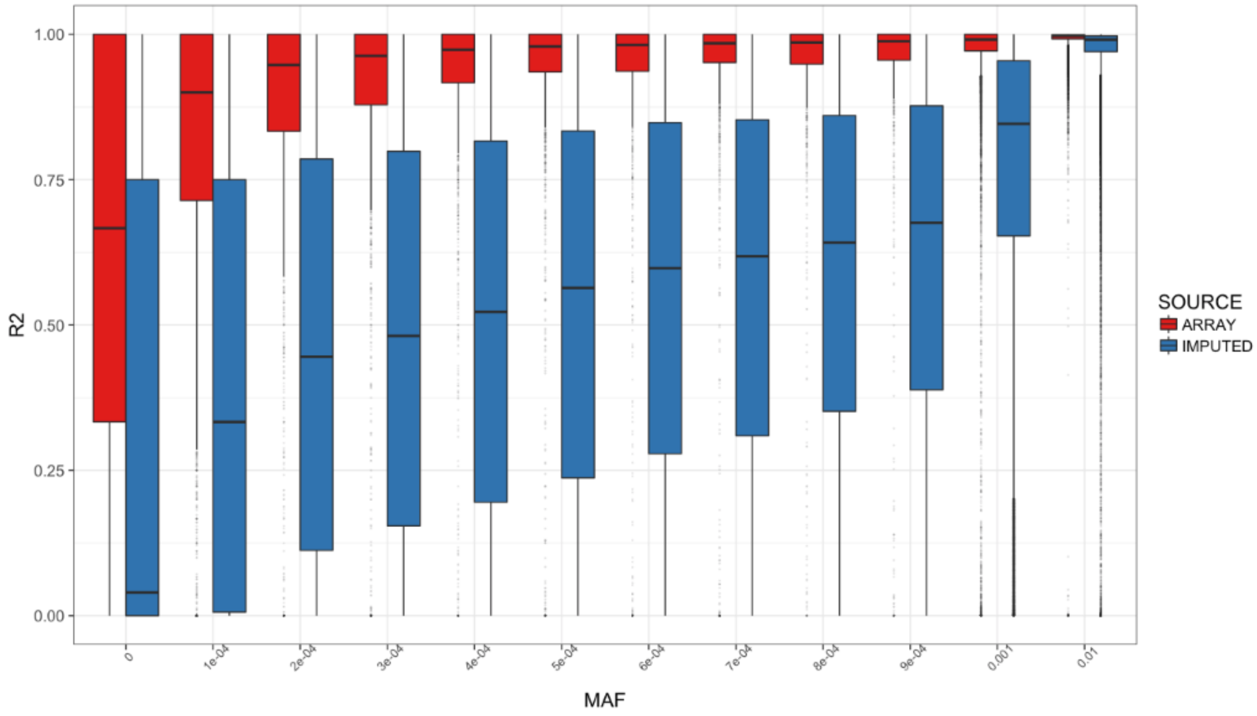


**Sup. Figure 3 | Concordance between WES-Array and WES-Imputed datasets**. R-squared correlation coefficients between variants in WES and imputed sequence (imputation score >0.3) (blue) and array genotypes and WES (red), calculated per variant and binned by minor allele frequency (MAF) in WES. Bin 0 includes variants MAF<$10^{-4}$. Bin 0.001 includes the range 0.001-0.01. Bin 0.01 includes the range 0.01-0.5. In total, n=46,912 individuals and n=75,334 variants were represented in the array-WES comparison, and n=46,860 individuals and n=899,455 variants with valid correlation coefficients were represented in the imputed-WES comparison.

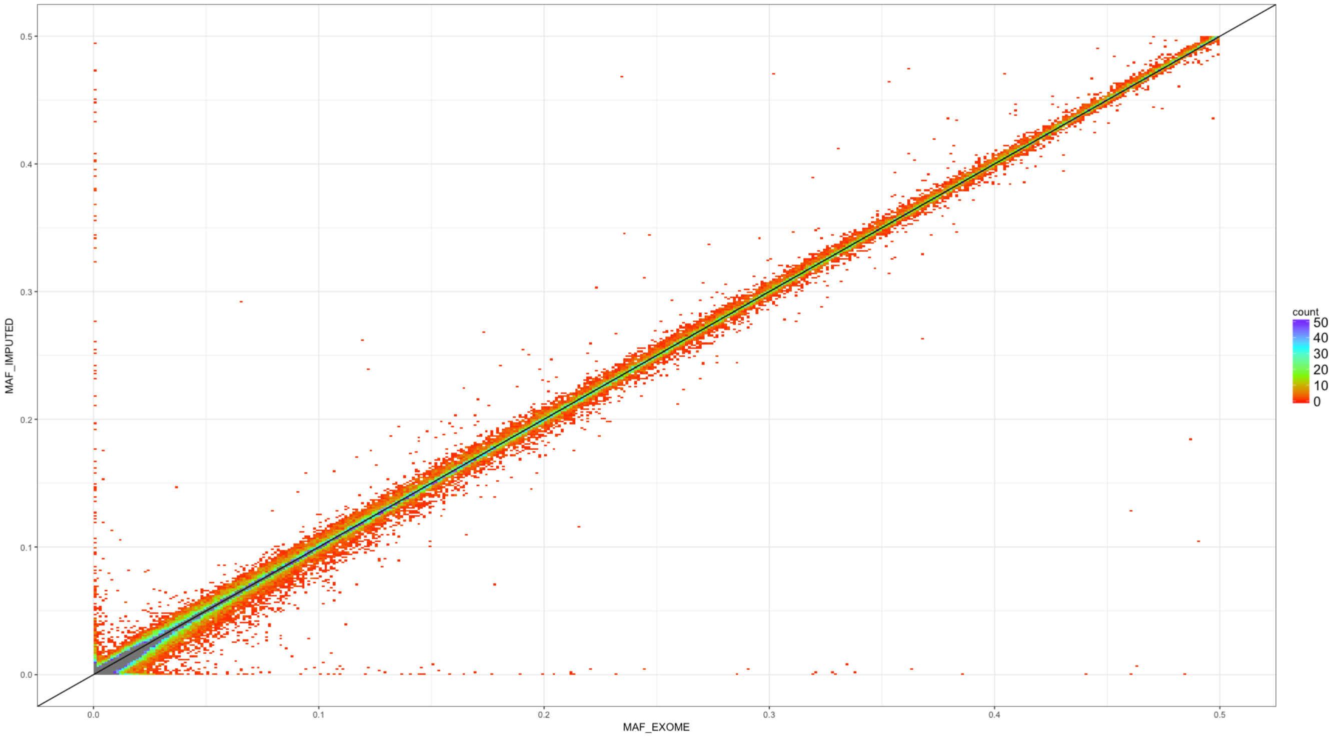


**<u>Sup. Figure 4 | Comparison of MAF in WES versus UKB imputed genotypes.</u>** n=1,065,159 autosomal variants in 46,860 European-ancestry individuals with both WES and imputed sequence (info score > 0.3) were included in the analysis. The black line shows x = y.

| | Variants in Imputed Sequence, info > 0.3, n=49,797 Participants | | Median Per Participant (IQR) | |
|---|---|---|---|---|
| | # Variants | # Variants MAF≤1% | # Variants | # Variants MAF≤1% |
| Total | 62,644,999 | 53,296,285 | 3,708,914 (18,537) | 39,395 (1,958) |
| Targeted Regions[1] | 780,445 | 708,506 | 26,129 (291) | 510 (46) |
| **Variant Type** [1] | | | | |
| SNVs | 767,266 | 697,707 | 25,220 (283) | 498 (45) |
| Indels | 13,179 | 10,799 | 909 (28) | 13 (5) |
| **Functional Prediction** | | | | |
| Synonymous | 254,423 | 226,453 | 10,615 (133) | 153 (22) |
| Missense | 396,021 | 367,467 | 9,681 (142) | 273 (29) |
| LOF (any transcript) | 18,055 | 17,012 | 340 (17) | 13 (5) |
| LOF (all transcripts) | 7,392 | 7,004 | 90 (11) | 6 (3) |

**Sup. Table 1 | Summary statistics for variants in imputed sequence**. Counts of autosomal variants observed in n=49,797 UKB participants with WES and imputed sequence by type/functional class for all and for MAF < 1% frequency. All variants have imputation info score > 0.3. Median count of variants and interquartile range (IQR) for all variants and for MAF < 1%. [1] Counts restricted to WES targeted regions.

| | AAF[1] | WES | Imputed 50k | Both |
|---|---|---|---|---|
| LOFs | All | 235,915 | 19,451 | 12,488 |
| | <0.01% | 220,261 | 9,325 | 5,093 |
| | 0.01-0.1% | 12,213 | 6,775 | 4,918 |
| | 0.1-1% | 2,242 | 2,062 | 1,628 |
| | 1-5% | 513 | 470 | 369 |
| | >5% | 686 | 819 | 480 |
| Synonymous | All | 1,241,804 | 267,479 | 216,983 |
| | <0.01% | 1,053,364 | 115,957 | 79,750 |
| | 0.01-0.1% | 125,315 | 89,558 | 78,348 |
| | 0.1-1% | 32,406 | 31,925 | 30,243 |
| | 1-5% | 9,909 | 9,424 | 9,114 |
| | >5% | 20,810 | 20,615 | 19,528 |
| Missense | All | 2,518,075 | 420,194 | 317,623 |
| | <0.01% | 2,237,837 | 205,690 | 133,023 |
| | 0.01-0.1% | 203,419 | 139,684 | 115,769 |
| | 0.1-1% | 44,454 | 43,969 | 39,807 |
| | 1-5% | 12,275 | 11,635 | 10,905 |
| | >5% | 20,090 | 19,216 | 18,119 |

**Sup. Table 2 | Ascertainment of variation in WES and imputed sequence**. Counts of autosomal predicted LOF, synonymous, and missense variation in 49,797 individuals with both WES and imputed sequence. No additional quality control beyond GL filtering for WES was applied, thus variant counts are greater than in Table 2. [1]Counts of variants in WES and Both datasets are binned by alt allele frequency (AAF) in WES. Counts in Imputed 50k are binned by AAF in WES if available, otherwise by AAF in imputed sequence.

| | | # Autosomal genes containing at least N LOFs, MAF < 1% | | | | | |
|---|---|---|---|---|---|---|---|
| **Zygosity** | **Genomic resource** | **1** | **5** | **10** | **25** | **50** | **100** |
| Het | 50k exome | 17,751 | 15,269 | 12,629 | 7,799 | 4,573 | 2,453 |
| | 50k imputed (all) | 9,108 | 7,082 | 5,994 | 4,315 | 2,978 | 1,915 |
| | 50k imputed (info > 0.3) | 8,763 | 6,999 | 5,933 | 4,297 | 2,965 | 1,911 |
| | 50k imputed (info > 0.5) | 7,501 | 6,237 | 5,413 | 4,040 | 2,821 | 1,848 |
| | 50k imputed (info > 0.8) | 4,053 | 3,430 | 3,088 | 2,355 | 1,829 | 1,339 |
| | 500k imputed (all) | 9,346 | 8,086 | 7,485 | 6,644 | 5,915 | 4,956 |
| | 500k imputed (info > 0.3) | 8,724 | 7,711 | 7,267 | 6,539 | 5,847 | 4,916 |
| | 500k imputed (info > 0.5) | 7,208 | 6,449 | 6,171 | 5,703 | 5,199 | 4,468 |
| | 500k imputed (info > 0.8) | 3,711 | 3,332 | 3,177 | 2,956 | 2,755 | 2,475 |
| Hom | 50k exome | 1,071 | 135 | 33 | 1 | 0 | 0 |
| | 50k imputed (all) | 789 | 74 | 7 | 0 | 0 | 0 |
| | 50k imputed (info > 0.3) | 789 | 74 | 7 | 0 | 0 | 0 |
| | 50k imputed (info > 0.5) | 770 | 74 | 7 | 0 | 0 | 0 |
| | 50k imputed (info > 0.8) | 627 | 58 | 7 | 0 | 0 | 0 |
| | 500k imputed (all) | 1,755 | 598 | 351 | 120 | 21 | 3 |
| | 500k imputed (info > 0.3) | 1,752 | 597 | 351 | 120 | 21 | 3 |
| | 500k imputed (info > 0.5) | 1,701 | 586 | 346 | 120 | 21 | 3 |
| | 500k imputed (info > 0.8) | 1,197 | 500 | 300 | 106 | 18 | 3 |

**Sup. Table 3 | Number of autosomal genes with heterozygous, homozygous LOF variants.** Count of genes with at least the specified number of LOFs (MAF < 1%) in UK Biobank participants of European ancestry in approximately 50k (n=46,827 with WES and imputed sequence) and 500k (462,427 individuals).

| Category | #Variants | % of Total Known ACMG59Variants | #Carriers | % of individuals with reportable variants |
|---|---|---|---|---|
| **Broad (P)** | 1209 | 3.8 | 3644 | 7.29 |
| **Broad + LP** | 197 | - | 251 | 0.50 |

**Sup. Table 4 | Variation in ACMG genes in UKB WES.** UKB participants are carriers of known pathogenic (P) and likely (LP) pathogenic variants in ACMG59 v2.0 genes based on broad variant filtering criteria. Notably, the number and proportion of individuals estimated to carry a reportable variant using the broad definition (See Supplemental Methods) is inconsistent with the population prevalence of Mendelian disorders.

| Category | Number of Variants | % of Total Known Variants | Number of Carriers | % of individuals with Reportable Variants |
|---|---|---|---|---|
| **Pathogenic (P)** | 540 | 7.31 | 1774 | 1.94 |
| **Likely Pathogenic (LP)** | 396 | - | 769 | 0.84 |
| **Total** | 936 | - | 2527 | 2.76 |

**Sup. Table 5 | Variation in ACMG genes in Regeneron-Geisinger DiscovEHR WES.** 2.76% of GHS participants are carriers of known pathogenic (P) and likely (LP) pathogenic variants in ACMG59 v2.0 genes based on the strict variant filtering criteria compared to 2.03% in UKB (See Supplemental Methods).

| Gene | ICD10 code, Binary Phenotype | Control RR|RA|AA | Case RR|RA|AA | OR (95% CI) | P-Value |
|---|---|---|---|---|---|
| ***BRCA1*** | Z40.0, Encounter for prophylactic surgery for risk factors related to malignant neoplasms | 43447|58|0 | 58|4|0 | 50.0 10.7, 233.0) | $9.3x10^{-7}$ |
| ***BRCA2*** | C61, Malignant neoplasm of prostate | 45282|986|2 | 594|31|0 | 2.4 (1.6, 3.5) | $3.8x10^{-5}$ |
| ***COL3A1*** | Z87.3, Personal history of diseases of the musculoskeletal system and connective tissue | 39147|5|0 | 39|1|0 | 198.2 (8.8, 4469.3) | $9.2x10^{-4}$ |
| ***COL4A4*** | R31, Hematuria | 44361|71|0 | 1314|12|0 | 0.4 (0.6, 0.3) | $1.6x10^{-6}$ |
| ***FLG*** | L30, Other and unspecified dermatitis | 45226|1465|6 | 87|6|0 | 2.1 (0.9, 4.6) | $7.4x10^{-2}$ |
| ***KRT5*** | L85.9, Epidermal thickening, unspecified | 45475|9|0 | 24|2|0 | 3.7 (2.1, 6.5) | $5.1x10^{-6}$ |
| ***LDLR*** | Z95.0, Presence of cardiac pacemaker | 40253|26|0 | 49|2|0 | 62.0 (7.1, 537.1) | $2.0x10^{-4}$ |
| ***LDLR*** | I25.1, Atherosclerotic heart disease of native coronary artery | 43787|28|0 | 1649|3|0 | 2.8 (1.1, 7.3) | $2.8x10^{-2}$ |
| ***PALB2*** | C50, Malignant neoplasm of breast | 45745|104|0 | 1102|9|0 | 3.6 1.7, 7.7) | $1.2x10^{-3}$ |
| ***RBM20*** | I34.0, Nonrheumatic mitral (valve) insufficiency | 44871|22|0 | 108|3|0 | 1.3 (1.1, 1.4) | $6.1x10^{-6}$ |
| ***SMAD6*** | M50, Cervical disc disorders | 45063|50|0 | 118|5|0 | 1.9 (1.5, 2.4) | $1.9x10^{-7}$ |

| Gene | Quantitative Phenotype | RR|RA|AA | Beta (95% CI) | P-Value |
|---|---|---|---|---|
| ***COL6A3*** | Corneal resistance factor mean | 35580|58|0 | -0.59 (-0.84,-0.34) | $4.7x10^{-6}$ |
| ***GP1BB*** | Platelet count | 45448|4|0 | -2.25 (-3.21,-1.30) | $3.7x10^{-6}$ |
| ***IL17RA*** | Monocyte count | 45395|19|0 | -1.13 (-1.58,-0.69) | $5.7x10^{-7}$ |
| ***JAK2*** | Platelet count | 45428|24|0 | 0.96 (0.57,1.35) | $1.4x10^{-6}$ |

| | | | | |
|---|---|---|---|---|
| ***RHAG*** | Mean sphered cell volume | 43469\|19\|0 | -1.07 (-1.51, -0.63) | $1.9 \times 10^{-6}$ |
| ***TET2*** | Eosinophil percentage | 44746\|179\|0 | -0.34 (-0.49, -0.20) | $4.3 \times 10^{-6}$ |
| ***TET2*** | Eosinophil count | 44745\|178\|0 | -0.34 (-0.48, -0.19) | $5.9 \times 10^{-6}$ |
| ***TMPRSS6*** | Mean corpuscular haemoglobin | 45333\|44\|0 | -0.73 (-1.02,-0.44) | $6.0 \times 10^{-7}$ |

**Sup. Table 6 | Extended list of positive controls.**

| Gene | UKB Phenotype | GHS Phenotype | OR (95% CI) | P-Value |
|---|---|---|---|---|
| ***PIEZO1*** | I83.9, Asymptomatic varicose veins of lower extremities | I83.9, Asymptomatic varicose veins of lower extremities | 3.8 (2.2,6.4) | $1.5x10^{-6}$ |

| Gene | UKB Phenotype | GHS Phenotype | Beta (95% CI) | P-Value |
|---|---|---|---|---|
| ***COL6A1*** | Corneal resistance factor mean | NA[1] | - | - |
| ***COL6A1*** | Corneal hysteresis mean | NA[1] | - | - |
| ***GMPR*** | Mean corpuscular haemoglobin | Mean corpuscular volume | 0.18 (0.09,0.27) | $8.0x10^{-5}$ |
| ***IQGAP2*** | Mean platelet thrombocyte volume | Mean platelet volume | 0.35 (0.21,0.49) | $7.8x10^{-7}$ |
| ***MEPE*** | Heel bone mineral density | Femoral neck BMD T-score | -0.19 (-0.48,0.11) | 0.22 |

**Sup. Table 7 | Replication of novel LOF burden associations in DiscovEHR.** [1]No comparable phenotype was available for testing in DiscovEHR.

| Single-point analysis | | | | | | | | | Drop one - RVT | |
|---|---|---|---|---|---|---|---|---|---|---|
| SNPID (rsID) | Ref | Alt | AAF | Beta | SE | P-Val | MAC | Functional Effect | P-Val LOO Burden | Delta P-Val Burden |
| 16:88727163:G:A | G | A | 6.9E-5 | 2.81 | 0.88 | 0.001 | 6 | stop gained | 1.06E-6 | 33.91 |
| 16:88721586:G:C | G | C | 1.1E-5 | 14.63 | 119.5 | 0.903 | 1 | stop gained | 2.63E-7 | 8.45 |
| 16:88716874:G:A | G | A | 1.1E-5 | 14.40 | 119.5 | 0.904 | 1 | stop gained | 2.59E-7 | 8.31 |
| 16:88738735:D:1 | TC | T | 1.1E-5 | 13.95 | 119.5 | 0.907 | 1 | frameshift | 2.45E-7 | 7.86 |
| 16:88721268:D:1 | CT | C | 1.1E-5 | 13.90 | 119.5 | 0.907 | 1 | frameshift | 2.43E-7 | 7.80 |
| 16:88726546:C:T | C | T | 1.1E-5 | 13.67 | 119.5 | 0.909 | 1 | splice donor | 2.34E-7 | 7.49 |
| 16:88723253:G:A (rs368895635) | G | A | 2.3E-5 | 3.77 | 1.42 | 0.008 | 2 | stop gained | 2.31E-7 | 7.42 |
| 16:88719588:G:A | G | A | 1.1E-5 | 13.57 | 119.5 | 0.910 | 1 | stop gained | 2.29E-7 | 7.35 |
| 16:88720229:C:A | C | A | 1.1E-5 | 13.55 | 119.5 | 0.910 | 1 | stop gained | 2.28E-7 | 7.32 |
| 16:88727072:D:1 | TC | T | 2.3E-5 | 3.59 | 1.42 | 0.011 | 2 | frameshift | 2.22E-7 | 7.12 |
| 16:88716359:A:G (rs776709730) | A | G | 2.3E-5 | 3.15 | 1.42 | 0.026 | 2 | splice donor | 1.94E-7 | 6.23 |
| 16:88736324:G:A | G | A | 4.6E-5 | 2.48 | 1.18 | 0.035 | 4 | stop gained | 1.67E-7 | 5.35 |

**Sup. Table 8 | Single-point and aggregate results for the 65 LOF variants in *PIEZO1*.** P-VAL Leave-One-Out (LOO) Burden is the p-value of the absence/presence test excluding the variant being tested. Delta P-Val Burden is the ratio of the p-value in the drop-one-out analysis compared to the burden test using all 65 variants. Burden summary statistics using all 65 variants in unrelated individuals: B=1.44, SE=0.26, p-value=3.12E-8, cMAF=0.00174, cMAC=152, cMAF cases=0.0066, cMAC cases=17, cMAF controls=0.0016, cMAC cases=135. Stepwise logistic regression selected 11/65 variants (AIC 11451). Burden summary statistics using the 11 variants: B=3.71, SE=0.437, p-value< 2e-16, cMAF=0.0003, cMAC=22, cMAF cases=0.005, cMAC cases=12, cMAF controls=0.0001, cMAC controls=10. When analysis is restricted to variants with MAC > 1, stepwise logistic regression selects 5/13 variants (AIC 11484). Burden summary statistics using the 5 variants: B=3.02, SE=0.53, p-value=8.92E-9.

| Variant | LD($R^2$) | 1 | 2 | 3 | 4 | 5 | 6 | 7 | 8 | 9 | 10 | **11** | **12** |
|---|---|---|---|---|---|---|---|---|---|---|---|---|---|
| 1 | 16:88716359:A:G | 1 | 1.1E-9 | 1.1E-9 | 1.1E-9 | 1.1E-9 | 2.1E-9 | 1.1E-9 | 2.1E-9 | 6.3E-9 | 4.2E-9 | 1.1E-9 | 2.1E-4 |
| 2 | 16:88716874:G:A | | 1 | 5.2E-10 | 5.2E-10 | 5.2E-10 | 1.1E-9 | 5.2E-10 | 1.1E-9 | 3.1E-9 | 2.1E-9 | 5.2E-10 | 2.1E-5 |
| 3 | 16:88719588:G:A | | | 1 | 5.2E-10 | 5.2E-10 | 1.1E-9 | 5.2E-10 | 1.1E-9 | 3.1E-9 | 2.1E-9 | 5.2E-10 | 8.1E-6 |
| 4 | 16:88721268:D:1 | | | | 1 | 5.2E-10 | 1.1E-9 | 5.2E-10 | 1.1E-9 | 3.1E-9 | 2.1E-9 | 5.2E-10 | 8.1E-6 |
| 5 | 16:88721586:G:C | | | | | 1 | 1.1E-9 | 5.2E-10 | 1.1E-9 | 3.1E-9 | 2.1E-9 | 5.2E-10 | 8.1E-6 |
| 6 | 16:88723253:G:A | | | | | | 1 | 1.1E-9 | 2.1E-9 | 6.3E-9 | 4.2E-9 | 1.1E-9 | 1.6E-5 |
| 7 | 16:88726546:C:T | | | | | | | 1 | 1.1E-9 | 3.1E-9 | 2.1E-9 | 5.2E-10 | 2.1E-5 |
| 8 | 16:88727072:D:1 | | | | | | | | 1 | 6.3E-9 | 4.2E-9 | 1.1E-9 | 1.6E-5 |
| 9 | 16:88727163:G:A | | | | | | | | | 1 | 1.3E-8 | 3.1E-9 | 2.8E-5 |
| 10 | 16:88736324:G:A | | | | | | | | | | 1 | 2.1E-9 | 8.6E-5 |
| 11 | 16:88738735:D:1 | | | | | | | | | | | 1 | 8.1E-6 |
| 12 | 16:88835545:G:A | | | | | | | | | | | | 1 |

**Sup. Table 9 | Linkage Disequilibrium (LD) Assessment for *PIEZO1*.** LD (r2) across the 11 LOF variants selected by stepwise regression for *PIEZO1* and a positive control (16:88835545 G A; highlighted in green) reported in the literature for PIEZO1. None of these 12 variants are in LD, $R^2$>0.01. When the burden test is adjusted for the previously reported variant rs2911463 (16:88835545 G A), the burden test p-value remains < 2E-16 (Akaike information criterion 11,444), which indicates that the burden is not tagging the reported variant.

| Gene | Phenotype | Exome 50k P | Imputed 50k P | Imputed 500k P |
|---|---|---|---|---|
| ***ASXL1*** | Platelet distribution width | $4.7x10^{-9}$ | NA[1] | NA[1] |
| ***ASXL1*** | Red blood cell erythrocyte distribution width | $2.4x10^{-11}$ | NA[1] | NA[1] |
| ***CALR*** | D47, Other neoplasms of uncertain behavior of lymphoid, hematopoietic and related tissue | $4.1x10^{-8}$ | NA[1] | NA[1] |
| ***CHEK2*** | Platelet crit | $7.9x10^{-8}$ | 0.039 | $2.0x10^{-3}$ |
| ***COL6A1*** | Corneal resistance factor mean | $3.6x10^{-10}$ | NA[1] | NA[1] |
| ***COL6A1*** | Corneal hysteresis mean | $2.1x10^{-8}$ | NA[1] | NA[1] |
| ***GMPR*** | Mean corpuscular haemoglobin | $1.1x10^{-8}$ | $6.2x10^{-6}$ | $4.3x10^{-34}$ |
| ***GP1BA*** | Mean platelet thrombocyte volume | $6.4x10^{-8}$ | NA[1] | NA[1] |
| ***HBB*** | Red blood cell erythrocyte distribution width | $5.8x10^{-8}$ | 0.76 | 0.21 |
| ***HBB*** | Red blood cell erythrocyte count | $1.7x10^{-9}$ | 0.23 | 0.32 |
| ***IL33*** | Eosinophil percentage | $5.4x10^{-12}$ | $1.4x10^{-11}$ | $7.5x10^{-83}$ |
| ***IL33*** | Eosinophil count | $3.3x10^{-10}$ | $7.6x10^{-9}$ | $3.9x10^{-71}$ |
| ***IQGAP2*** | Mean platelet thrombocyte volume | $1.1x10^{-19}$ | 0.10 | $2.2x10^{-7}$* |
| ***KALRN*** | Mean platelet thrombocyte volume | $2.7x10^{-23}$ | $1.9x10^{-20}$ | $2.0x10^{-95}$ * |
| ***KLF1*** | Red blood cell erythrocyte distribution width | $1.5x10^{-13}$ | 0.43 | 0.72 |
| ***KLF1*** | Mean corpuscular haemoglobin | $1.7x10^{-16}$ | 0.77 | 0.036 |
| ***KLF1*** | Mean corpuscular volume | $4.0x10^{-14}$ | 0.63 | 0.028 |
| ***MEPE*** | Heel BMD | $1.4x10^{-8}$ | $5.9x10^{-3}$ | $1.6x10^{-14}$ |
| ***MLH1*** | Z85.0, Personal history of malignant neoplasm of digestive organs | $3.5x10^{-11}$ | NA[1] | 0.79 |
| ***PIEZO1*** | I83.9, Asymptomatic varicose veins of lower extremities | $2.7x10^{-8}$ | 0.084 | 0.27 |
| ***PKD1*** | N18, Chronic kidney disease | $2.9x10^{-10}$ | NA[1] | NA[1] |
| ***TTN*** | I42, Cardiomyopathy | $1.4x10^{-8}$ | 0.015 | 0.052 |
| ***TUBB1*** | Platelet distribution width | $2.5x10^{-23}$ | $5.4x10^{-3}$ | $1.0x10^{-21}$ |
| ***TUBB1*** | Mean platelet thrombocyte volume | $2.4x10^{-8}$ | 0.028 | $4.8x10^{-10}$* |
| ***TUBB1*** | Platelet count | $2.1x10^{-9}$ | 0.029 | $1.6x10^{-8}$ |

**Sup. Table 10 | Comparison of LOF burden associations in 50k exome vs 50k imputed vs 500k imputed.** *Result computed using linear regression. [1]No LOFs were available for testing in imputed data.

| Single-point Analysis | | | | | | | | | Drop one - RVT | |
|---|---|---|---|---|---|---|---|---|---|---|
| **SNPID** | **REF** | **ALT** | **AAF** | **BETA** | **SE** | **P-Val** | **MAC** | **Functional Effect** | **P-Val LOO Burden** | **Delta P-Val Burden** |
| 4:87834767:D:4 | CAGTA | C | 3.7E-5 | 0.56 | 0.57 | 3.3E-1 | 3 | Splice region | 1.29E-5 | 1.56 |
| 4:87838631:G:A | G | A | 9.9E-5 | -0.27 | 0.35 | 4.4E-1 | 8 | Splice acceptor | 1.71E-6 | 0.21 |
| 4:87839684:G:A | G | A | 1.2E-5 | -0.26 | 0.99 | 8.0E-1 | 1 | Splice acceptor | 6.90E-6 | 0.84 |
| 4:87839693:C:G | C | G | 1.2E-5 | -0.12 | 0.99 | 9.0E-1 | 1 | Stop gained | 7.28E-6 | 0.88 |
| 4:87844983:D:1*$ | GA | G | 2.5E-5 | 2.12 | 0.70 | 2.5E-3 | 2 | frameshift | 3.72E-5 | 4.51 |
| 4:87845066:D:4*$ | GGAAA | G | 4.5E-4 | 0.51 | 0.17 | 2.0E-3 | 36 | frameshift | 8.00E-4 | 97.01 |
| 4:87845210:G:A | G | A | 1.2E-5 | 1.19 | 0.99 | 2.3E-1 | 1 | start_lost | 1.23E-5 | 1.49 |
| 4:87845320:I:7* | T | TATATGGG | 1.2E-5 | 1.69 | 0.99 | 8.9E-2 | 1 | frameshift | 1.49E-5 | 1.81 |
| 4:87845359:I:1 | T | TG | 7.4E-5 | 0.42 | 0.40 | 3.0E-1 | 6 | frameshift | 1.45E-5 | 1.76 |
| 4:87845484:D:1*$ | AG | A | 9.0E-4 | 0.35 | 0.12 | 2.7E-3 | 72 | frameshift | 9.53E-4 | 115.46 |
| 4:87845585:I:1 | C | CG | 1.2E-5 | -0.64 | 0.99 | 5.2E-1 | 1 | frameshift | 5.90E-6 | 0.72 |
| 4:87845726:D:1 | AG | A | 1.2E-5 | -0.22 | 0.99 | 8.2E-1 | 1 | frameshift | 7.01E-6 | 0.85 |
| 4:87845732:D:4* | AAGTG | A | 1.2E-5 | 1.49 | 0.99 | 1.3E-1 | 1 | frameshift | 8.25E-6 | 1.00 |
| 4:87845741:I:5 | T | TGACAA | 1.2E-5 | 1.49 | 0.99 | 1.3E-1 | 1 | frameshift | 8.25E-6 | 1.00 |
| 4:87845761:D:1 | CA | C | 1.2E-5 | 1.17 | 0.99 | 2.4E-1 | 1 | frameshift | 1.22E-5 | 1.48 |
| 4:87846011:D:1 | GA | G | 5.0E-5 | -0.53 | 0.50 | 2.9E-1 | 4 | frameshift | 2.51E-6 | 0.30 |

**Sup. Table 11 | Single-point and aggregate results for 16 LOF variants in MEPE.** Burden summary statistics using the 16 variants in unrelated subjects: beta= -0.375, se=0.084, p-value=8.39E-6, cumulative minor allele frequency (cMAF)=0.00174, cumulative minor allele count (cMAC)=140. None of these variants individually account for the whole aggregate signal, as illustrated by the drop-one-out analyses. Stepwise regression selected 5/16 variants (AIC -380.96). Burden summary statistics using the 5 variants: beta=-0.456, se=0.094, p-value=1.14E-6, cMAF=0.0014, cMAC=112. When we restrict analysis to variants with MAC > 1, stepwise logistic regression selects 3/7 variants (AIC -379.59). Burden summary statistics using the 3 variants: beta=-0.436, se=0.095, p-value=4.14E-6. * Variants selected by stepwise regression. $ Variants selected by stepwise regression when the full model contains variants with MAC > 1.

| | | Exome Sequencing Variants | | | | | Previously reported variants | | | | | |
|---|---|---|---|---|---|---|---|---|---|---|---|---|
| **Variant** | **LD (r2)** | **1** | **2** | **3** | **4** | **5** | **6** | **7** | **8** | **9** | **10** | **11** |
| **1** | 4:87844983:D:1 | 1 | 4.2E-8 | 1.1E-9 | 8.2E-8 | 1.1E-9 | 8.4E-6 | 6.6E-8 | 9.3E-6 | 6.3E-8 | 6.2E-8 | 2.1E-6 |
| **2** | 4:87845066:D:4 | | 1 | 2.1E-8 | 1.6E-6 | 2.1E-8 | 2.0E-4 | 5.1E-4 | 2.7E-3 | 5.0E-1 | 1.2E-6 | 1.2E-4 |
| **3** | 4:87845320:I:7 | | | 1 | 4.1E-8 | 5.2E-10 | 3.1E-5 | 4.9E-5 | 7.4E-6 | 3.2E-8 | 3.1E-8 | 2.2E-5 |
| **4** | 4:87845484:D:1 | | | | 1 | 4.1E-8 | 3.4E-8 | 8.3E-4 | 3.0E-4 | 2.5E-6 | 7.8E-1 | 3.9E-4 |
| **5** | 4:87845732:D:4 | | | | | 1 | 2.0E-6 | 3.3E-8 | 7.4E-6 | 3.2E-8 | 3.1E-8 | 6.9E-6 |
| **6** | 4:87055207:A:T | | | | | | 1 | 4.5E-4 | 1.9E-3 | 2.7E-5 | 9.1E-7 | 1.4E-3 |
| **7** | 4:87724192:C:T | | | | | | | 1 | 4.0E-3 | 5.5E-4 | 6.8E-4 | 7.7E-4 |
| **8** | 4:87832908:C:T | | | | | | | | 1 | 4.9E-3 | 2.1E-4 | 3.1E-2 |
| **9** | 4:87882603:A:G | | | | | | | | | 1 | 1.9E-6 | 3.2E-4 |
| **10** | 4:87887987:T:C | | | | | | | | | | 1 | 3.4E-4 |
| **11** | 4:87910097:G:A | | | | | | | | | | | 1 |

**Sup. Table 12 | LD (r2) across the 5 LOF variants contributing to the MEPE burden signal and the six previously reported variants.** *MEPE* is a known locus for heel BMD with 6 conditionally independent signals within or close to it[2]. Two of the 6 variants are in partial (r2=0.5) or high (r2=0.78) LD with two of the variants contributing to the burden test (green). To determine whether the burden result is explained by any of these previously reported variants, we recalculated the burden analysis after conditioning on each of the 6 previously reported variants separately. We observe that the burden cannot be fully explained by the single-point previously reported variants (maximum p-value is 3.6x10-6), while the burden p-value increases to 2x10-4 when we condition jointly on all six. We conclude that our burden result strengthens supporting evidence of real association for *MEPE*, points to LoF variants likely to be driving the signal (none of the six positive controls are known to be functional).

| Single-point Analysis | | | | | | | | | Drop one - RVT | |
|---|---|---|---|---|---|---|---|---|---|---|
| SNPID | REF | ALT | AAF | BETA | SE | P-Val | MAC | Functional Effect | P-Val LOO Burden | Delta P-Val Burden |
| 21:45981876:D:17*$ | CCCTGCTGC TGCAGGCCT | C | 3.08E-5 | -1.73 | 0.71 | 1.5E-2 | 2 | frameshift | 1.48E-6 | 19.26 |
| 21:45981944:C:T*$ | C | T | 6.16E-5 | -1.43 | 0.50 | 4.3E-3 | 4 | stop gained | 5.35E-6 | 69.60 |
| 21:45984285:C:T | C | T | 1.54E-5 | -0.70 | 1.00 | 4.8E-1 | 1 | stop gained | 7.64E-8 | 0.99 |
| 21:45989090:C:T* | C | T | 1.54E-5 | -2.11 | 1.00 | 3.5E-2 | 1 | stop gained | 5.76E-7 | 7.49 |
| 21:45989138:G:A | G | A | 1.54E-5 | -1.05 | 1.00 | 2.9E-1 | 1 | splice donor | 1.29E-7 | 1.67 |
| 21:45990977:A:G* | A | G | 1.54E-5 | -1.93 | 1.00 | 5.4E-2 | 1 | splice acceptor | 4.48E-7 | 5.83 |
| 21:45990995:D:1* | CT | C | 1.54E-5 | -1.40 | 1.00 | 1.6E-1 | 1 | frameshift | 2.12E-7 | 2.75 |
| 21:45992184:I:1 | G | GC | 1.54E-5 | -0.98 | 1.00 | 3.3E-1 | 1 | frameshift | 1.15E-7 | 1.50 |
| 21:45998172:G:A | G | A | 1.54E-5 | -2.27 | 1.00 | 2.3E-2 | 1 | splice donor | 7.19E-7 | 9.35 |
| 21:45998172:I:1 | G | GT | 1.54E-5 | -1.10 | 1.00 | 2.7E-1 | 1 | splice region | 1.38E-7 | 1.79 |
| 21:45999683:D:1 | TG | T | 1.54E-5 | -0.09 | 1.00 | 9.3E-1 | 1 | frameshift | 3.06E-8 | 0.40 |

**Sup. Table 13 | Single-point and aggregate results for *COL6A1* with Corneal hysteresis.** Single-point and aggregate results for 11 LOF variants in COL6A1. Burden summary statistics using the 11 variants: beta= -1.388, se=0.258, p-value=7.69E-8, cMAF=0.0002, cMAC=15. None of these variants individually account for the whole aggregate signal, as illustrated by the drop-one analyses. Stepwise regression selected 5/11 variants (AIC -17.68). Burden summary statistics using the 5 variants: beta= -1.623, se=0.333, p-value=1.12E-6, cMAF=0.00014, cMAC=9. When we restrict analysis to variants with MAC > 1, stepwise regression selects both those variants (AIC -13.33). Burden summary statistics using the 2 variants: beta= -1.529, se=0.408, p-value=0.00018. * Variants selected by stepwise regression. $ Variants selected by stepwise regression when the full model contains variants with MAC > 1.

| Single-point Analysis | | | | | | | | | Drop one - RVT | |
|---|---|---|---|---|---|---|---|---|---|---|
| SNPID | REF | ALT | AAF | BETA | SE | PV | MAC | Functional Effect | P-Val LOO Burden | Delta P-Val Burden |
| 21:45981876:D:17*$ | CCCTGCTGC TGCAGGCCT | C | 3.08E-5 | -1.95 | 0.71 | 5.8E-3 | 2 | frameshift | 4.43E-7 | 39.21 |
| 21:45981944:C:T*$ | C | T | 6.16E-5 | -1.36 | 0.50 | 6.5E-3 | 4 | stop gained | 5.02E-7 | 44.39 |
| 21:45984285:C:T | C | T | 1.54E-5 | -0.69 | 1.00 | 4.9E-1 | 1 | stop gained | 1.03E-8 | 0.91 |
| 21:45989090:C:T* | C | T | 1.54E-5 | -2.58 | 1.00 | 9.8E-3 | 1 | stop gained | 1.80E-7 | 15.94 |
| 21:45989138:G:A* | G | A | 1.54E-5 | -1.48 | 1.00 | 1.4E-1 | 1 | splice donor | 3.50E-8 | 3.10 |
| 21:45990977:A:G* | A | G | 1.54E-5 | -2.10 | 1.00 | 3.5E-2 | 1 | splice acceptor | 8.95E-8 | 7.92 |
| 21:45990995:D:1 | CT | C | 1.54E-5 | -1.29 | 1.00 | 2.0E-1 | 1 | frameshift | 2.63E-8 | 2.33 |
| 21:45992184:I:1 | G | GC | 1.54E-5 | -0.85 | 1.00 | 4.0E-1 | 1 | frameshift | 1.32E-8 | 1.17 |
| 21:45998172:G:A | G | A | 1.54E-5 | -2.85 | 1.00 | 4.4E-3 | 1 | splice donor | 2.63E-7 | 23.31 |
| 21:45998172:I:1 | G | GT | 1.54E-5 | -0.24 | 1.00 | 8.1E-1 | 1 | splice region | 5.02E-9 | 0.44 |
| 21:45999683:D:1 | TG | T | 1.54E-5 | -0.68 | 1.00 | 5.0E-1 | 1 | frameshift | 1.02E-8 | 0.90 |

**Sup. Table 14 | Single-point and aggregate results for *COL6A1* with Corneal resistance.** Single-point and aggregate results for 11 LOF variants in COL6A1. Burden summary statistics using the 11 variants: beta= -1.474, se=0.258, p-value=1.13E-8, cMAF=0.0002, cMAC=15. None of these variants individually account for the whole aggregate signal, as illustrated by the drop-one analyses. Stepwise regression selected 5/11 variants (AIC -23.07). Burden summary statistics using the 5 variants: beta= -1.724, se=0.333, p-value=2.32E-7, cMAF=0.00014, cMAC=9. When we restrict analysis to variants with MAC > 1, stepwise regression selects both those variants (AIC -15.58). Burden summary statistics using the 2 variants: beta= -1.558, se=0.408, p-value=0.000136. * Variants selected by stepwise regression. $ Variants selected by stepwise regression when the full model contains variants with MAC > 1.

| | | Exome Sequencing Variants | | | | | Previously Reported Variants | | | | | | | | | | |
|---|---|---|---|---|---|---|---|---|---|---|---|---|---|---|---|---|---|
| **Variant** | **LD (r2)** | **1** | **2** | **3** | **4** | **5** | **6** | **7** | **8** | **9** | **10** | **11** | **12** | **13** | **14** | **15** | **16** |
| **1** | 21:45981876:D:17 | 1 | 5.2E-9 | 1.0E-9 | 1.0E-9 | 1.0E-9 | 8.2E-7 | 1.9E-6 | 2.3E-6 | 2.4E-7 | 1.1E-6 | 2.0E-6 | 1.2E-6 | 7.7E-5 | 3.4E-5 | 2.9E-6 | 3.2E |
| **2** | 21:45981944:C:T | | 1 | 2.6E-9 | 2.6E-9 | 2.6E-9 | 2.0E-6 | 4.7E-6 | 1.4E-5 | 3.0E-4 | 9.9E-5 | 1.5E-5 | 3.1E-6 | 1.1E-4 | 5.7E-5 | 3.7E-5 | 9.5E |
| **3** | 21:45989090:C:T | | | 1 | 5.2E-10 | 5.2E-10 | 4.1E-7 | 9.3E-7 | 1.2E-6 | 1.4E-5 | 2.0E-5 | 1.0E-6 | 6.2E-7 | 3.3E-7 | 5.0E-7 | 1.5E-6 | 9.9E |
| **4** | 21:45990977:A:G | | | | 1 | 5.2E-10 | 4.1E-7 | 5.4E-4 | 1.2E-6 | 2.0E-5 | 2.0E-5 | 6.2E-5 | NA | 3.8E-5 | 5.6E-5 | 1.5E-6 | 9.9E |
| **5** | 21:45990995:D:1 | | | | | 1 | 4.1E-7 | 9.3E-7 | 1.2E-6 | 2.0E-5 | 2.0E-5 | 1.0E-6 | 6.2E-7 | 3.3E-7 | 5.0E-7 | 1.5E-6 | 9.9E |
| **6** | 21:45512700_C_A | | | | | | 1 | 3.5E-5 | 1.8E-4 | 8.6E-5 | 1.1E-3 | 3.2E-4 | 2.8E-6 | 2.3E-3 | 2.2E-3 | 7.4E-5 | 2.3E |
| **7** | 21:45734434_C_A | | | | | | | 1 | 2.0E-2 | 3.3E-3 | 7.8E-3 | 1.9E-2 | 6.4E-1 | 1.2E-2 | 9.5E-3 | 3.1E-5 | 1.6E |
| **8** | 21:45906833_A_G | | | | | | | | 1 | 2.1E-1 | 2.9E-1 | 8.7E-1 | 1.7E-2 | 6.0E-2 | 2.7E-1 | 8.5E-5 | 8.9E |
| **9** | 21:45909076_T_G | | | | | | | | | 1 | 1.2E-1 | 1.9E-1 | 3.4E-3 | 9.0E-2 | 5.9E-2 | 1.1E-5 | 1.1E |
| **10** | 21:45952033_G_A | | | | | | | | | | 1 | 3.1E-1 | 5.3E-3 | 7.1E-3 | 7.0E-2 | 1.1E-4 | 2.2E |
| **11** | 21:45957382_G_A | | | | | | | | | | | 1 | 1.7E-2 | 7.2E-2 | 2.9E-1 | 1.6E-4 | 7.6E |
| **12** | 21:45984508_C_G | | | | | | | | | | | | 1 | 1.1E-2 | 1.6E-2 | 3.7E-6 | 1.1E |
| **13** | 21:46002498_C_T | | | | | | | | | | | | | 1 | 8.3E-2 | 7.7E-4 | 4.9E |
| **14** | 21:46022420_C_T | | | | | | | | | | | | | | 1 | 1.5E-4 | 1.2E |
| **15** | 21:46099621_T_C | | | | | | | | | | | | | | | 1 | 1.5E |
| **16** | 21:46542898_A_G | | | | | | | | | | | | | | | | 1 |

**Sup. Table 15 | LD (r2) across the LOF variants contributing to the *COL6A1* burden signal and the 11 previously reported variants.** While some positive controls are in partial LD with each other, the variants contributing to the burden are not in LD with any of the previously reported variants.

| Variant | Study | Phenotype | RR\|RA\|AA | Beta (95% CI) | P |
|---|---|---|---|---|---|
| rs753138805 | UKB 500k imputed | Heel bone mineral density | 409177\|377\|0 | -0.41 (-0.50,-0.32) | $8.1x10^{-19}$ |
| rs753138805 | HUNT | BMD | 19401\|303\|1 | -0.53 (-0.65,-0.41) | $2.1x10^{-18}$ |

| Variant | Study | Phenotype | RR\|RA\|AA | OR (95% CI) | P |
|---|---|---|---|---|---|
| rs753138805 | UKB 500k imputed | M81, Osteoporosis without current pathological fracture | Ctrl:452235\|406\|0 Case:3478\|6\|0 | 1.9 (0.9,4.2) | 0.10 |
| rs753138805 | HUNT | Any Fracture | Ctrl:44936\|543\|1 Case:23753\|402\|0 | 1.4 (1.2,1.5) | $1.6x10^{-5}$ |
| rs753138805 | HUNT | Fracture of ankle and foot | Ctrl:44936\|543\|1 Case:5368\|110\|0 | 1.8 (1.4,2.4) | $3.3x10^{-6}$ |
| rs753138805 | HUNT | Fracture of hand or wrist | Ctrl:44936\|543\|1 Case:5761\|102\|0 | 1.5 (1.2,1.9) | $7.8x10^{-4}$ |
| rs753138805 | HUNT | Fracture of upper limb | Ctrl:44936\|543\|1 Case:10927\|201\|0 | 1.5 (1.3,1.8) | $1.2x10^{-5}$ |
| rs753138805 | HUNT | Fracture of unspecified bones | Ctrl:44936\|543\|1 Case:8480\|147\|0 | 1.5 (1.2,1.8) | $3.4x10^{-4}$ |
| rs753138805 | HUNT | Fracture of lower limb | Ctrl:44936\|543\|1 Case:7732\|133\|0 | 1.4 (1.1,1.8) | $1.7x10^{-3}$ |
| rs753138805 | HUNT | Fracture of pelvis | Ctrl:44936\|543\|1 Case:799\|17\|0 | 1.9 (1.0,3.5) | 0.043 |
| rs753138805 | HUNT | Fracture of vertebral column without mention of spinal cord injury | Ctrl:44936\|543\|1 Case:1990\|33\|0 | 1.4 (0.9,2.0) | 0.14 |
| rs753138805 | HUNT | Torus fracture | Ctrl:43203\|519\|1 Case:805\|13\|0 | 1.4 (0.7,2.7) | 0.30 |
| rs753138805 | HUNT | Fracture of ribs | Ctrl:44936\|543\|1 Case:1635\|25\|0 | 1.2 (0.9,1.9) | 0.38 |
| rs753138805 | HUNT | Skull and face fracture and other intercranial injury | Ctrl:65020\|889\|1 Case:1922\|29\|0 | 1.1 (0.8,1.7) | 0.58 |

**Sup. Table 16 | Results of *MEPE* rs753138805 associations in UKB 500k imputed and Nord-Trøndelag Health Study (HUNT).** The first of two *MEPE* LOFs with the most significant single variant associations with BMD, rs753138805 (p-value = $1.4x10^{-3}$), encodes a four base-pair deletion that leads to an early truncation. We tested this variant (Imputation $R^2$ = 0.71) for association with BMD and osteoporosis in all European-descent UKB participants with imputed sequence and phenotype of interest available. Replication of the rs753138805 association with BMD and extension to fractures was completed in HUNT (Imputation $R^2$ = 0.99). rs778732516, which encodes a single base-pair deletion in *MEPE* (BMD p-value =$6.2x10^{-5}$) was not present in the UKB imputed sequence nor HUNT. Effect size for BMD measures are in standard deviations from the mean. Effect size for fractures and osteoporosis are odds ratios.
