## Supplemental Methods for "Whole exome sequencing and characterization of coding variation in 49,960 individuals in the UK Biobank"

#### WES sample preparation and sequencing

Genomic DNA samples normalized to approximately 16 ng/ul were transferred to the Regeneron Genetics Center from the UK Biobank in 0.5ml 2D matrix tubes (Thermo Fisher Scientific) and stored in an automated sample biobank (LiCONiC Instruments) at -80°C prior to sample preparation. One sample had insufficient DNA for sequencing. Exome capture was completed using a high-throughput, fully-automated approach developed at the Regeneron Genetics Center. Briefly, DNA libraries were created by enzymatically shearing 100ng of genomic DNA to a mean fragment size of 200 base pairs using a custom NEBNext Ultra II FS DNA library prep kit (New England Biolabs) and a common Y-shaped adapter (Integrated DNA Technologies) was ligated to all DNA libraries. Unique, asymmetric 10 base pair barcodes were added to the DNA fragment during library amplification with KAPA HiFi polymerase (KAPA Biosystems) to facilitate multiplexed exome capture and sequencing. Equal amounts of sample were pooled prior to overnight exome capture, approximately 16 hours, with a slightly modified version of IDT's xGen probe library; supplemental probes were added to capture regions of the genome well-covered by a previous capture reagent (NimbleGen VCRome) but poorly covered by the standard xGen probes (design bed file available by request). In total, n=38,997,831 bases were included in the targeted regions. Captured fragments were bound to streptavidin-coupled Dynabeads (Thermo Fisher Scientific) and non-specific DNA fragments removed through a series of stringent washes using the xGen Hybridization and Wash kit according to the manufacturer's recommended protocol (Integrated DNA Technologies). The captured DNA was PCR amplified with KAPA HiFi and quantified by qPCR with a KAPA Library Quantification Kit (KAPA Biosystems). The multiplexed samples were pooled and then sequenced using

75 base pair paired-end reads with two 10 base pair index reads on the Illumina NovaSeq 6000 platform using S2 flow cells.

### Sequence alignment, variant identification, and genotype assignment

Upon completion of sequencing, raw data from each Illumina NovaSeq run was gathered in local buffer storage and uploaded to the DNAnexus platform[1] for automated analysis. After upload was complete, analysis began with the conversion of CBCL files to FASTQ-formatted reads and assigned, via specific barcodes, to samples using the bcl2fastq conversion software (Illumina Inc., San Diego, CA). Sample-specific FASTQ files, representing all the reads generated for that sample, were then aligned to the GRCh38 genome reference with BWA-mem[2]. The resultant binary alignment file (BAM) for each sample contained the mapped reads' genomic coordinates, quality information, and the degree to which a particular read differed from the reference at its mapped location. Aligned reads in the BAM file were then evaluated to identify and flag duplicate reads with the Picard[3] MarkDuplicates tool, producing an alignment file (duplicatesMarked.BAM) with all potential duplicate reads marked for exclusion in downstream analyses.

GVCF files, including variant calls, were then produced on each individual sample using the WeCall variant caller[4], identifying both SNVs and INDELs as compared to the reference. Additionally, each GVCF file carried the zygosity of each variant, read counts of both reference & alternate alleles, genotype quality representing the confidence of the genotype call, and the overall quality of the variant call at that position.

Upon completion of variant calling, individual sample BAM files were converted to fully lossless CRAM files using samtools[5]. Metric statistics were captured for each sample to evaluate capture, alignment, insert size, and variant calling quality, using Picard[3], bcftools[6], and FastQC[7].

Following completion of sample sequencing, samples showing disagreement between genetically-determined and reported sex (n=15), high rates of heterozygosity/contamination (D-stat > 0.4) (n=7), low sequence coverage (less than 85% of targeted bases achieving 20X coverage) (n=1), or geneticallyidentified sample duplicates (n=14), and WES variants discordant with genotyping chip (n=9) were excluded. Six samples failed quality control in multiple categories, resulting in 38 individuals being excluded. The remaining 49,960 samples were then used to compile a project-level VCF (PVCF) for downstream analysis. The PVCF was created using the GLnexus joint genotyping tool[8]. Care was taken to carry all homozygous reference, heterozygous, homozygous alternate, and no-call genotypes into the project-level VCF. An additional filtered PVCF, '*Goldilocks*' (GL), was also generated. In the filtered GL PVCF, any SNV genotype with read depth less than seven reads (DP < 7) was changed to a no-call. After the application of the DP genotype filter, only SNV variant sites that met at least one of the following two criteria were retained: 1) at least one heterozygous variant genotype with allele balance ratio greater than or equal to 15% (AB >= 0.15); 2) at least one homozygous variant genotype. The same filtering was applied to INDEL variants but with an INDEL depth filter of DP < 10 and an INDEL allele balance cutoff of AB >= 0.20. Multi-allelic variant sites in the PVCF file were normalized by left-alignment and represented as bi-allelic.

### Phenotype definition

ICD10-based cases required one or more of the following: a primary diagnosis or ≥ 2 secondary diagnosis in in-patient Health Episode Statistics (HES) records. ICD10-based excludes had ≥1 primary or ≥2 secondary diagnosis in the code range. ICD10-based controls were defined as those individuals that were not cases or excluded. Custom phenotype definitions included one or more of the following: ICD-10 diagnosis, self-reported illness from verbal interview and physician-diagnosed illness from online-follow-up, touchscreen information. Quantitative measures (e.g. physical measures, blood counts, cognitive function tests, imaging derived phenotypes) were downloaded from UKB repository and spanned one or more visits. In total, data for 3,390 field IDs were downloaded from UKB repository. We selected ~1,225 field IDs for WES association tests based on sample size and preliminary genetic utility. These field IDs expanded to 1,073 binary traits with case count ≥50 and 669 quantitative traits for testing as dependent variables in WES association analyses. Lists of binary and quantitative field IDs are included in extended data ExtData_TraitLists.xlsx.

#### Extrapolation of genes with LOF variant carriers in 500k

To estimate the number of genes with heterozygous LOF carriers in WES in 500k individuals, we counted the number of genes with N LOF carriers in sample size X as an estimate of the number of genes with 10N LOF carriers in 10X (Sup. Methods Table 1, Sup. Methods Fig. 1).

| # Genes | N, Minimum number of carriers of heterozygous LOF variants in 50k | N, Predicted minimum number of carriers of heterozygous LOF variants in 500k |
|---|---|---|
| 17,751 | 1 | 10 |
| 15,269 | 5 | 50 |
| 12,629 | 10 | 100 |
| 7,799 | 25 | 250 |
| 4,573 | 50 | 500 |
| 2,453 | 100 | 1,000 |
| 1,022 | 250 | 2,500 |
| 432 | 500 | 5,000 |
| 64 | 1,000 | 10,000 |

| # Genes | N, Minimum number of carriers of heterozygous LOF variants in 15k | N, Predicted minimum number of carriers of heterozygous LOF variants in 150k |
|---|---|---|
| 16,449 | 1 | 10 |
| 10,661 | 5 | 50 |
| 6,864 | 10 | 100 |
| 3,102 | 25 | 250 |
| 1,636 | 50 | 500 |
| 810 | 100 | 1,000 |
| 177 | 250 | 2,500 |
| 13 | 500 | 5,000 |
| 1 | 1,000 | 10,000 |

**Sup. Methods Table 1 | Predicted number of genes with heterozygous LOF carriers in larger WES sample sizes from existing 50k WES data.** The number of autosomal genes with at least 1, 5, 10, etc. heterozygous LOF carriers passing Goldilocks QC and genotype missingness<10%, and HWE p-value>$10^{-15}$ in 46,808 UKB participants of European ancestry with WES, and predicted number of genes with N heterozygous LOF carriers in 150k and 500k.

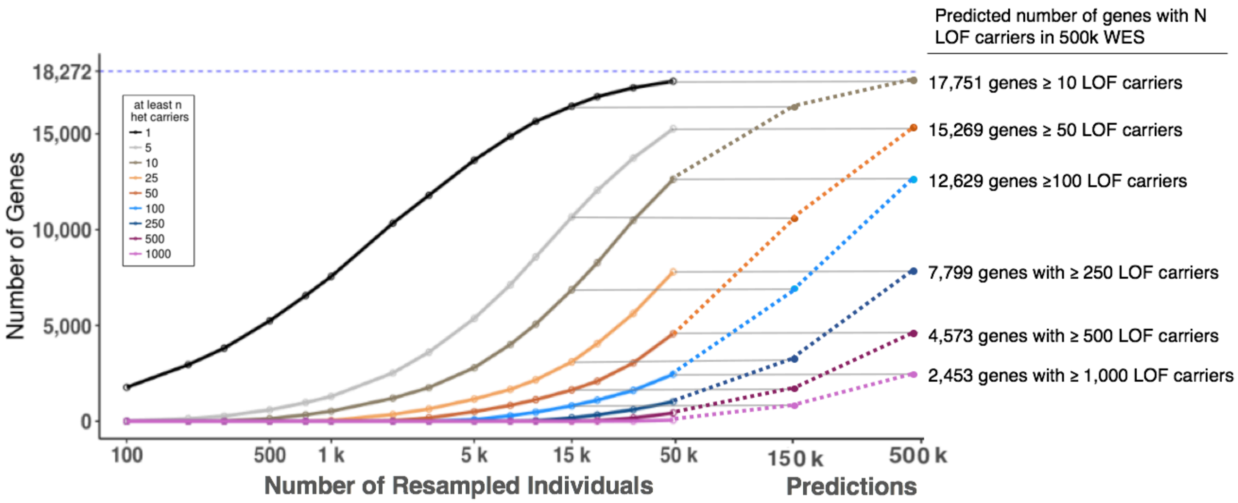


**Sup. Methods Figure 1 | Predicted number of genes with heterozygous LOF carriers in ~500k WES from existing WES data.** The number of autosomal genes with at least 1, 5, 10, etc. heterozygous and LOFs passing Goldilocks QC and genotype missingness<10%, and HWE p-value>$10^{-15}$ increases with sample size. 46,808 UKB participants of European ancestry with WES were down-sampled at random to the number of individuals specified on the horizontal axis. The number of genes containing at least the indicated count of LOF carriers MAF<1% as in the legend are plotted on the vertical axis. The maximum number of autosomal genes is 18,272 in this analysis. Horizontal lines connect the number of genes with at least N heterozygous LOF carriers in 50k to the predicted number of heterozygous LOF carriers in 500k.

### Annotation of predicted loss-of-function (LOF) variants

We annotated variants using SnpEff[9] and gene models from Ensembl[10] Release 85. We obtained a comprehensive and high-quality transcript set for protein coding regions which included all protein coding transcripts with an annotated Start and Stop codon from the Ensembl gene models. Variants annotated as stop_gained, start_lost, splice_donor, splice_acceptor, stop_lost and frameshift are considered predicted LOF variants.

A recent large-scale study of genetic variation in 141,456 individuals, gnomAD, provides a catalog of LOF variants[11]. A direct comparison to this data is difficult due to numerous factors such as differences in exome sequencing capture platforms, variant calling algorithms, annotation and number of individuals. Additionally, the geographic distribution of ascertainment (and thus genetic diversity) in the Non-Finnish Europeans (NFE) subset of gnomAD may be larger than that of UK Biobank with WES in this report. Nevertheless, we annotated the gnomAD exome sites labeled as "PASS" from gnomAD r2.1 using our annotation pipeline (Sup. Methods Table 1). Data from gnomAD were lifted over to HG38 using Picard LiftoverVcf. We obtain 514,325 LOFs in the autosomes of 125,748 exomes. In comparison, 515,326 LOFs are reported in gnomAD including exome sequence and 15,708 genomes. Further, we subset the gnomAD data to NFE restricted to variants with $MAF_{NFE} < 1\%$ (Supplemental Methods Table 1). LOFs were also subset to those restricted to canonical transcripts. The canonical transcripts were defined using the Ensembl definition (http://useast.ensembl.org/Help/Glossary?id=346).

We repeated similar calculations with a set of "high confidence" LOFs as defined by LOFTEE[11]. The total number of LOFs differ depending on the transcript models used for annotation. As expected, the number of variants that lead to LOF consequence in all transcripts of a gene are fewer than the number of LOF variants that affect any transcript of a gene (Sup. Methods Table 1).

| **LOF filtering and transcript definition** | **UKB WES,** n=46,979 | | **gnomAD,** n=56,885 | |
|---|---|---|---|---|
| | **#LOFs** | **#Genes** | **#LOFs** | **#Genes** |
| LOFs in any transcript | 202,127 | 17,751 | 260,431 | 17,946 |
| LOFs in canonical transcript | 189,430 | 17,624 | 244,251 | 17,825 |
| LOFs in all transcripts | 134,693 | 16,130 | 174,731 | 16,458 |
| LOFTEE filtered LOFs in any transcript | 189,519 | 17,640 | 244,669 | 17,856 |
| LOFTEE filtered LOFs in canonical transcript | 179,357 | 17,510 | 231,486 | 17,732 |
| LOFTEE filtered LOFs in all transcripts | 128,222 | 16,006 | 166,321 | 16,349 |

**Sup. Methods Table 2 | LOF variants in UKB and gnomAD using RGC annotation pipeline.** Number of autosomal predicted loss of function (LOF) variants MAF <1% and number of genes with at least one heterozygous LOF in UKB participants of European ancestry, n=46,979 and gnomAD Non-Finnish-Europeans, n=56,885. MAF was estimated and applied within UKB and gnomAD samples, respectively. LOF in "any" transcript is a variant that is annotated as a LOF variant in at least one transcript of a gene. LOF in "all" transcripts is a variant that affects all transcripts of a gene and has a predicted LOF consequence

### Visual validation of variants in genes with 1 heterozygous LOF variant

We know that singleton variants which are the only predicted LOF variant seen in a given gene will be enriched for false positive calls. Thus, to improve the accuracy in our estimates of the number of LOF carrying genes, we performed visual validation on all 684 singletons that were the only LOF variants seen in their respective genes.

We interrogated these single-sample LOF variants within IGV[12] by reviewing the CRAM read stacks. Various criteria can be used to make determinations of variant status (here we simply classified variants as either likely real or likely false) and are subjective depending on the reviewer. Characteristics available for review (described below) include, but are not limited to, mapping quality (MQ), depth (DP), allele balance (AB), read tiling, read strandedness, variant position within reads, duplicate read status, insert size, and mate pair consensus.

- MQ - While an MQ of 60 is best, there are many real variants called in low mappability regions. So, variants with a preponderance of MQ 0 reads are considered to be likely false, as are variants without at least one MQ 60 read supporting the call. Variants are generally considered suspect where there is little read support with unique alignments, tools like BLAT[13] can be used to confirm unique alignments for questionable reads.

- DP – Low-end depth cutoffs for our filtered variants are 7 reads for SNPs and 10 reads for INDELs. However, reviewers are cautious of sites with significant read depth, where depth may be due to homologous or repetitive sequence and will likely have a lower MQ reflecting secondary mapping sites. This can be validated by viewing the region in UCSC's Genome Browser's[14] available RepeatMasker track, but generally sites at the extremes of coverage are considered suspect.
- AB - In our Goldilocks filtered data, AB is at a site-level and is only one-sided. AB cutoffs for SNPs is 15/85 and 20/80 for INDELs. Variants outside this cutoff are considered suspect, and poor allele balance is strongly associated with likely false variants.
- Read Tiling/Coverage – This is different than depth. There ought to be even and consistent tiling of reads across the targets within the capture design. Variants that fall in regions with poor tiling often come at the edges of target regions, and are likely to have issues with read strand bias.
- Read strandedness – We treat sites with a bias to one read direction as suspect. Again, using caution at edges of targets where the data is more prone to reads from only one strand, but strandedness at the edge of a capture target should be consistent between alt and ref allele reads, and properly overlap the target region.
- Variant Position – The ends of reads are error prone, so variants only seen in the ends of reads are suspect.
- Duplicate Read Status – Variants whose evidence comes primarily from reads sharing the same or nearly the same start and end points (particularly in low MAPQ regions) are considered suspect.
- Insert Size – The insert size distribution in this data is quite consistent, and generally should be consistent with templates on the order of 140-240bp long. Variants supported by read-pairs with wildly different or inconsistent insert size are considered suspect.
- Mate Pair Consensus – Where a read's mate pair overlaps, both reads should have the same sequence, and variants with evidence of inconsistent mate-pair consensus are suspect.

While visual validation is an inherently subjective process, visual validation using these criteria has been shown to correlate will with Sanger validation. For the review of this data two analysts worked independently and cross-validated a variety of the most questionable sites. Extended data ExtData_SingletonpLOFVizVal.xlsx indicates which of the singleton LOF variants were reviewed (n=685), with likely false variants (n=106) denoted with VV=0, comprising 15.5% of the variants that were reviewed. As expected, the validation rate for SNPs was higher than that of INDELs, with 241 of the 292 INDEL variants reviewed validated (82.5%) and 338 of 392 SNPs reviewed validated (86.2%).

When performing visual validation on homozygous reference exome calls (as with the comparison between discordant imputed calls and exome data), the reads were examined for any evidence of variation by looking for any reads supporting the called allele, for low quality alternate allele bases, and for mis-aligned, multi-nucleotide polymorphism, or soft-clipped reads containing alternate alleles at the location of the variant call. Evidence of the called allele were accounted for separately from evidence of any alternate allele as the former represents a possible undercall in the exome, while the latter is likely a misrepresentation of the variation in the region by either the imputation, exome, or both.

### Concordance between WES, array and imputed genotypes

We calculated concordance as follows: we computed the squared correlation between the allele dosages for one datatype (WES) versus the other (imputed hard calls/array genotypes) for one variant at a time. For average estimates, R-squared values were pooled across bins of allele frequency defined by the exome dataset[15,16].

### Methods for ACMG59 medically actionable variant survey

We compiled two lists of known pathogenic (P) variants reported in the ClinVar and HGMD (Human Gene Mutation Database) databases in all the ACMG59 recommended genes (Sup. Methods Figure 2):

1. A high-confidence (Strict set) conservative list of 316 variants classified as "Pathogenic" or "Likely Pathogenic" with no conflicting interpretations based on stringent review and assertion criteria for

clinical significance (≥2stars) was assembled from the NCBI ClinVar database. The full ClinVar dataset of variants and their corresponding classifications was downloaded from: ftp://ftp.ncbi.nlm.nih.gov/pub/clinvar/vcf_GRCh38/archive_2.0/2018/clinvar_20180429.vcf.gz

2. A comprehensive list of 1,209 pathogenic variants (Broad set) compiled from variants reported in the HGMD and ClinVar databases. This set is a union of all high-confidence disease-causing "DM" variants from HGMD (2017-12-19 version) and ClinVar "Pathogenic" and "Likely Pathogenic" without conflicting interpretations variants. From this set, variants that had discordant pathogenic annotations between ClinVar and HGMD were removed.

In addition to pathogenic variants, for both the Strict and Broad sets, we also included LOF variants in the 45 genes where, according to the ACMG recommendations[17,18], protein truncation is known to be the disease-causing mechanism and are therefore classified as likely pathogenic (LP) variants. Thus, this category also includes predicted LOF variants that may be present in HGMD but not reported in ClinVar and Pathogenic/Likely Pathogenic LOF variants in ClinVar that did not meet the assertion criteria (≥2 stars) for the Strict set of variants.

A total of 608 candidate medically actionable variants were identified within the Strict P + LP dataset. Of these, 50 variants were flagged as failing the allele balance threshold of at least 20% of reads calling the alternate allele in at least one sample. All variants in the Strict dataset were visually inspected by expert curators using the Integrative Genomics Viewer (IGV). Thirteen of the flagged variants (13/50) were confirmed to be of high quality albeit with allele balance ratios between 15-19%. Additionally, 14 variants were deemed "unclear" with good quality but few (2-3) reads only calling the alternate allele; whereas 33 variants failed visual confirmation likely representing sequence and/or alignment errors with several occurring in the vicinity of poly-A tracts and low complexity regions. Of note, rs587779193 [hg38.chr2:47414419(G>T)] a 3 star "Likely Pathogenic" splicing variant in *MSH2* reported in ClinVar was excluded from the dataset as it occurred in the vicinity of a flagged indel variant adjacent to a low-complexity region that is prone to misalignment and consequently a likely error. All other 33 failed variants were LP variants of which 22 were indels. The final list of 555 "Strict" known pathogenic (P, n=316) and likely pathogenic (LP, n=239) variants identified in the 49,960 UKB participants with WES is included as Extended Data (ExtData_ACMGVariants_V2.xlsx). This set affects 1,000 non-redundant individuals (2.0% of cohort). About 58% of the variants identified were single nucleotide polymorphisms (N=323), whereas 41% were indels (N=226). Additionally, we identified and visually validated 10 complex alleles, 6 of which were dinucleotide substitutions and 4 were complex indel variants in *cis*.

For the "Broad" category, we made a union dataset of pathogenic and high confidence disease-causing variants from ClinVar and HGMD respectively along with the LP LOF set in the ACMG59 genes. 420 variants labelled "Benign"/"Likely Benign" or with conflicting interpretations of pathogenicity in ClinVar, or annotated as low confidence in HGMD, were removed. This resulted in 1,406 candidate medically actionable variants in 3,876 carriers (7.76% of cohort). Of note, of the 239 LP variants included in the Strict set, 47 of them were found in the Broad dataset, meaning that they have been reported in HGMD but not in ClinVar or that they do not meet the stringent $\leq$2 star ClinVar classification that we used for defining the variant datasets. Overall, the higher estimate of potentially actionable findings using a "Broad" set of pathogenic variants partly reflects the increased number of "disease-causing" variants reported in HGMD that have been proven to be not pathogenic upon reassessment and identification in a large number of unaffected carriers[19-22].

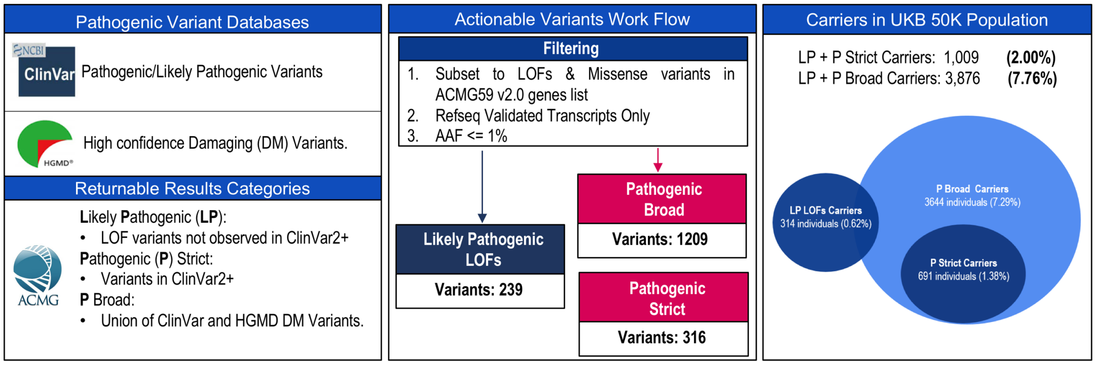


**Sup. Methods Figure 2 | Actionable Variants Work Flow**

### Definition of UK Biobank phenotypes

| Characteristics | Definition |
|---|---|
| Townsend deprivation index | Field ID: 189; measure of material deprivation within a population |
| Inpatient ICD10 codes per patient | Field IDs: 41142 (primary diagnosis), 41078 (secondary diagnosis)<br>1 or more ICD 3D primary or secondary diagnoses in HES data |
| Percentage of patients with >=1 ICD10 diagnoses | Field IDs: 41142 (primary diagnosis), 41078 (secondary diagnosis)<br>Percentage of patients with 1 or more primary or secondary diagnoses (ICD 3D) in HES data |
| **Cardiometabolic phenotypes** | **ICD10 codes, self-reported and doctor-diagnosed code** |
| Coronary Disease | I20 OR I21 I22 I23 OR I24 OR I25 OR 42000_0__0_Date_of_first_myocardial_infarction_CASE (!=NA) OR Z951 OR Z955 |
| Heart Failure | I50 |
| Type 2 Diabetes | E11 |
| **Respiratory and immunological phenotypes** | |
| Asthma | J45 OR J46 OR Self_reported_asthma (1111) OR Doctor_diagnosed_asthma |
| COPD | J41 OR J42 OR J43 OR J44 |
| Rheumatoid Arthritis | M05 OR M06 OR Self-reported_Rheumatoid_Arthritis (1464) |
| Inflammatory Bowel Disease | K50 OR K51 OR Self-reported_Ulceritive_Colitis (1463) |
| **Neurodegenerative phenotypes** | |
| Alzheimer's Disease | G30 |
| Parkinson's Disease | G20 |
| Multiple Sclerosis | G35 |
| Myasthenia Gravis | G70 |
| **Oncology phenotypes** | |
| Breast cancer | HES(C50) OR Self-Reported_Breast_Cancer (1002) OR Cancer Registry (C50) |
| Ovarian cancer | HES(C56) OR Self-Reported_Ovarian_Cancer (1039) OR Cancer Registry (C56) |
| Prostate cancer | HES(C61) OR Self-Reported_Prostate_Cancer (1026) OR Cancer Registry (C61) |
| Pancreatic cancer | HES(C25) OR Self-Reported_Pancreatic_Cancer (1044) OR Cancer Registry (C50) |

| | |
|---|---|
| Melanoma | HES(C43) OR Self-Reported_Melanoma (1049) OR Cancer Registry (C43) |
| Lung cancer | HES(C34) OR Self-Reported_Lung_Cancer (1001) OR Cancer Registry (C34) |
| Colorectal cancer | HES(C18) OR Self-Reported_Colorectal_Cancer (1049) OR Cancer Registry (C18) |
| Cutaneous squamous cell carcinoma (CSCC) | HES(C44) OR Self-Reported_Squamous_Cell_Carcinoma (1062) OR Cancer Registry (C44) |
| **Enhanced measures** | **Field ID** |
| Pulse rate | 4194 Instance 0 |
| Visual Acuity Measured | 5187 Instance 0 |
| IOP measured (left) | 5262 Instance 0 |
| Autorefraction | 5159 Instance 0 |
| Retinal OCT | 6072 Instance 0 |
| ECG at rest | 22334 |
| Cognitive Function | 20242, Fluid intelligence completion status |
| Digestive Health | 21066, Born by caesarian section |
| Physical Activity Measurement | 90112, Fraction acceleration <= 25 milli-gravities |

**Sup. Methods Table 3 | Definition of UK Biobank phenotypes**. Disease phenotype cases were defined using >=1 primary or >=2 secondary diagnoses.

### Methods for LOF Burden Association Analysis

We performed burden tests of association for rare predicted LOFs within 49,960 individuals of European ancestry with WES. For each gene region as defined by Ensembl[10]. LOFs with MAF ≤ 0.01 were collapsed such that any individual that is heterozygous for at least one LOF in that gene region is considered heterozygous, and only individuals that carry two copies of the same LOF are considered homozygous. We did not phase rare variants, and so compound heterozygotes are not considered in this analysis. We additionally performed LOF burden analysis in the full 500k imputed dataset as well as a 50k imputed dataset down-sampled to the same set of individuals with WES. For these data, imputed LOFs were first converted to hardcalls using PLINK v2.0 (no imputation quality threshold, alternate allele dosage must be within 0.1 of nearest hardcall to be non-missing) and then collapsed exactly as described for WES.

For each gene region, 668 rank-based inverse normal transformed (RINT) quantitative measures (including all subjects and sex-stratified models) with ≥5 individuals with non-missing phenotype information were assessed using an additive mixed model implemented in BOLT-LMM v2[23]. Prior to normalization, traits were first transformed as appropriate (log10, square) and adjusted for a standard set of covariates including age, sex, study site, first four principal components of ancestry, and in some cases BMI and/or smoking status. Data-points greater than five median absolute deviations from the median were excluded as outliers prior to normalization. 1,073 discrete outcomes (including all subjects and sex-stratified models) with ≥ 50 cases were assessed with covariate adjustment for age, sex and first four principle components of ancestry using a generalized mixed model implemented in SAIGE[24]. For each quantitative and discrete trait included in the analysis, only gene regions in which > 3 LOF carriers with non-missing phenotype and covariate information were evaluated.

We systematically defined positive controls using a two-step approach. First, we annotated each gene for relevant disease, trait, biological, or functional evidence using publicly available resources including OMIM[25], NCBI MedGen, and the NHGRI-EBI GWAS catalogue[26]. For those genes with supporting evidence from at least one source, we then manually curated NCBI PubMed to verify the relationship between the trait and LOF variants in the gene of interest. Genes with locus-level support for the trait of interest or related phenotype(s) in the GWAS catalog but lacking clear supporting evidence for a LOF association are reported herein as novel LOF associations.

### Methods for single variant LOF Association Analysis

We performed single variant association analysis using the same methods as described in the methods section for burden association analysis. For gene-trait associations with $p < 10^{-7}$ (Tables 5 and 6), we calculated single variant association statistics with the phenotype of interest for all LOFs included in the burden test that are observed with a minor allele count ≥ 5 in the 49,960 European ancestry individuals with WES. Association statistics for these variants are reported in Extended Data (ExtData_SingleVariantLOFs_V1.xlsx).

### Methods for Replication and Follow-up

For all novel LOF associations (Table 6), we aimed to replicate the observed association in European ancestry individuals of the Regeneron-Geisinger DiscovEHR study[27]. Gene-sets for LOF burden analysis were created as previously described. For replication of quantitative traits associations, analysis was completed using linear regression with covariate adjustment for age, age-squared, sex, and first four principal components of ancestry implements in PLINK v1.9[28]. For replication of discrete outcomes associations, analysis was completed using Firth-penalized logistic regression with covariate adjustment for age, age-squared, sex, and first four principal components of ancestry implements in PLINK v1.9. All analyses in DiscovEHR were completed in two separate batches (60k and 30k data freezes) and subsequently meta-analyzed using PLINK v1.9.

For follow-up of *MEPE* rsXXX, we leveraged data from the full UK Biobank 500k imputed dataset and the Nord-Trøndelag Health Study (HUNT)[29] (Sup. Methods Table 3). Analyses in the 500k imputed dataset were performed as described in "Methods for LOF Burden Association Analysis." All analyses in HUNT was completed using SAIGE.

| **Phenotype** | **ICD9** | **ICD10** |
|---|---|---|
| Fracture of lower limb | 820-823;825;827;905;V54 | S72;S82;S92;T02;T12 |
| Fracture of ankle and foot | 824-826 | S82;S92 |
| Fracture of pelvis | 808 | S32 |
| Fracture of upper limb | 810-813;818-819;905;V54 | S22;S42;S52;T02;T10;T92 |
| Fracture of hand or wrist | 814-817 | S52;S62 |
| Fracture of vertebral column without mention of spinal cord injury | 805-806;905;V54 | M49;S12;S22;S32;T02;T91 |
| Fracture of ribs | 807 | S22 |
| Fracture of unspecified bones | 807;809;828-829;905;V54;V66-67 | S22;S32;T02;Z09;Z54 |

| | | |
|---|---|---|
| Skull and face fracture and other intercranial injury | 800-804;854;905;907;V15 | S02;S06;T02;T06;T90 |
| Torus fracture | 823 | |

**Sup. Methods Table 3 | Definition of HUNT fracture phenotypes**. Fracture phenotype cases were defined using ICD9 and ICD10 codes.
